# Complementary biological and computational approaches identify distinct mechanisms of chlorpyrifos versus chlorpyrifos oxon induced dopaminergic neurotoxicity

**DOI:** 10.1101/2022.07.15.500261

**Authors:** Shreesh Raj Sammi, Tauqeerunnisa Syeda, Kendra D. Conrow, Maxwell C. K. Leung, Jason R. Cannon

## Abstract

Organophosphate (OP) pesticides are widely used in agriculture. While acute cholinergic toxicity has been extensively studied, chronic effects on other neurons are less understood. Here, we demonstrated that the OP pesticide chlorpyrifos (CPF) and its oxon metabolite are dopaminergic neurotoxicants in *Caenorhabditis elegans*. CPF treatment led to inhibition of mitochondrial complex II, II + III, and V in rat liver mitochondria, while CPF oxon did not (complex II + III, and IV inhibition observed only at high doses). While the effect on *C. elegans* cholinergic behavior was mostly reversible with toxicant washout, dopamine-associated deficits persisted, suggesting dopaminergic neurotoxicity was irreversible. CPF reduced the mitochondrial content in a dose-dependent manner and the fat modulatory genes *cyp-35A2* and *cyp-35A3* were found to have a key role in CPF neurotoxicity. These findings were consistent with *in vitro* effects of CPF and CPF oxon on nuclear receptor signaling and fatty acid/steroid metabolism observed in ToxCast assays. Two-way hierarchical analysis revealed *in vitro* effects on estrogen receptor (ER,) pregnane X receptor (PXR), and peroxisome proliferator-activated receptor gamma (PPAR gamma) pathways as well as neurotoxicity of chlorpyrifos, malathion, and diazinon, while these effects were not detected in malaoxon and diazoxon. Taken together, our study suggests that mitochondrial toxicity and metabolic effects of CPF, but not CPF-oxon, have a key role of CPF neurotoxicity in the low-dose, chronic exposure. Further mechanistic studies are needed to examine mitochondria as a common target for all OP pesticide parent compounds, since this has important implications on cumulative pesticide risk assessment.

## 1. Introduction

Organophosphate (OP) pesticides are an important class of agricultural chemicals with a global sale of USD $7 billion in 2017 (Morder Intelliegence 2022). In the U.S.A., OPs account for approximately 30% of all insecticide usage from 2012 to 2016 (Morder Intelliegence 2022; U.S. Environmental Protection Agency (US EPA) 2017). The usage of OP pesticides has been in decline in recent years, partly due to ecological and human health concerns (Costa et al. 2008a; Landis et al. 2020; Richardson et al. 2019) as well as the introduction of newer pesticide classes such as pyrethroids and neonicotinoids (U.S. Environmental Protection Agency (US EPA) 2017). Yet, OP pesticides remain a popular option for pest management among corn, cotton, and specialty crop farmers. Chlorpyrifos (CPF) – an OP pesticide that raised public concern over its adverse neurological effects in humans – was banned for all agricultural uses by California Environmental Protection Agency in 2019 (Calfornia Department of Pesticide Regulation (CDPR) 2018; California Environmental Protection Agency (Cal EPA) 2019). Yet, other OP pesticides – including ethephon, malathion, and naled – collectively accounted for over 1,000,000 pounds of pesticide usage in California in 2018 (as compared to 600,000 pounds for CPF alone)(California Department of Pesticide Regulation (CDPR)). Since OP pesticides are still used in large quantities in the U.S.A. and other parts of world, there is a dire need to understand their mechanism of toxicity as a major class of environmental pollutants.

Cholinesterase inhibition is considered by the U.S. Environmental Protection Agency (U.S. EPA) to be a critical effect to assess the human health hazards following exposure to OP pesticides (U.S. Environmental Protection Agency (U.S. EPA) 2000). It is also considered the primary mechanism by which many insecticide OPs induce neurotoxicity in humans, which is a key assumption to justify dose additivity in OP cumulative risk assessment (U.S. Environmental Protection Agency (US EPA) 2006). With the toxicity primarily linked to acetylcholine (Ach) neurons, effect on other neuron types is relatively understudied. Recently, studies on Swiss albino mice have demonstrated a possible link between CPF and PD(Ali et al. 2019); Emerging evidence suggests that cholinesterase inhibition is not the sole contributor to OP insecticide neurotoxicity. Using the U.S. EPA ToxCast database, we demonstrated that much of the OP bioactivity interfered with cytochrome P450 and lipid/steroid metabolism rather than cholinergic signaling (Leung and Meyer 2019). The peroxisome proliferator-activated receptor (PPAR) pathway, a key nuclear receptor regulatory pathway for lipid/steroid metabolism, was found to respond to chlorpyrifos exposure *in vitro* in a dose-dependent manner (Herriage et al. 2022). Additionally, other neuron types such as dopamine (DA) neurons are also vulnerable to chlorpyrifos exposure (Eddins et al. 2010; Singh et al. 2018; Zhang et al. 2015). Yet, it remains unclear whether an alternative common mechanism exists to account for the non-cholinergic effects of OP exposure.

While recent data suggests potential effects on mitochondrial enzymes (Turton et al. 2021), detailed information across broad dose ranges and active metabolites of CPF/OP pesticides is still obscure. The aim of this study is to identify novel primary mechanisms of non-cholinesterase neurotoxicity using complementary biological and computational approaches. We examined the connection between dopaminergic neurotoxicity, mitochondrial toxicity, and fat metabolism in the *Caenorhabditis elegans* model. This was followed by testing the effect of CPF and its active metabolite, CPF Oxon on dopaminergic neurotoxicity, DA associated behavior vis a vis acetylcholine associated behavior, and mitochondrial damage and physiological deficits. Next, we mined the U.S. EPA ToxCast database to profile the *in vitro* effect of chlorpyrifos, malathion, diazinon, and their oxon metabolites on nuclear receptor signaling, cytochrome P450, and other enzymes involved in lipid and steroid metabolism. These results are expected to advance understanding of non-acetylcholine OP neuronal targets, along with possible links to neurological disease.

## 2. Material and Methods

### 2.1. Culture and Maintenance of *C. elegans* Strains

*C. elegans* strains, Bristol N2, BZ555, RB2046, VC710 and *Escherichia coli* OP50, were procured from *Caenorhabditis* Genetics Centre, (University of Minnesota, Minnesota) and grown on nematode growth medium (NGM) at 22 °C. Age-synchronization was achieved by sodium hypochlorite method followed by overnight incubation in M9 buffer.

### 2.2. Treatment with CPF and CPF Oxon

100 mM stocks of CPF and CPF oxon were prepared in ethanol. L1 stage worms were treated with different exposure levels of CPF and CPF oxon and in liquid media as described (Boyd et al. 2012). For CPF: doses ranging from 0 to 25 μM, 0 to 500 μM, and 0 to 150 μM were used for behavioral, neurodegeneration, and mitochondrial complex assay respectively. For CPF oxon: doses ranging from 0 to 5 μM, 0 to 500 μM, and 0 to 150 μM were used for behavioral, neurodegeneration, and mitochondrial complex assay respectively. In case of washout experiments, worms were treated for two days and allowed to recover for 1 day before the assays.

### 2.3. *C. elegans* Neurodegeneration Assay

Neuropathological effects of CPF and CPF oxon on dopaminergic neurons, was studied by treating L1 stage worms with different concentrations (0 to 500 μM) for 72 hrs at 22 °C. Neurodegeneration was quantified as described by Yao et al. (2010) (Yao et al. 2010) with slight modifications. Briefly, treated worms were washed three times using M9 buffer and anesthetized using 10 μL of 100 mM sodium azide. Counting of neurons was done for all neuron types i.e., Cepahlic sensila (CEP), Anterior deirid (ADE) and posterior deirid (PDE) using FITC filter. *C. elegans* has eight DA neurons, 4 CEP, 2 ADE, and 2 PDE (Sammi et al. 2018b). The percentage of intact neurons (PIN) was calculated for a minimum of 20 worms per group (independent replicates).

### 2.4. Mitotracker staining and quantification

MitoTracker staining was performed as described (Gaffney et al. 2014), with slight modifications. MitoTracker Red CM-H2Xros is the reduced form of mitotracker which specifically stains the viable mitochondria (referred to as mitochondrial content) based on membrane potential (Invitrogen 2008). Briefly, MitoTracker Red CM-H2Xros was added to liquid culture on day 2 and mixed gently. MitoTracker Red (final concentration, 4.72 μM) was fed to worms by mixing it with *E. coli* OP50. Worms were washed using M9 buffer on day 3, incubated with *E. coli* OP50 solution for 30 min, and washed again three times. Worms were anesthetized using 10 μL of 100 mM sodium azide and observed using TX red filter. Approximately 20 worms per group were analyzed semi-quantitatively using Image J (Schneider et al. 2012), excluding the gut region from ROI.

### 2.5. Assay for Mitochondrial Enzyme Activity

Assessment of mitochondrial complex activity for complex I to IV was conducted as described by Spinazzi et al., 2012 (Spinazzi et al. 2012). Briefly, mitochondria were isolated from rat liver and treated with different doses of CPF and CPF oxon for 15 minutes. The substrate was added and absorbance was recorded every 10 seconds for 3 minutes. Relative enzyme activity (EA) was calculated as: (Δ Absorbance/min × 1,000)/[(extinction coefficient × volume of sample used in ml) × (sample protein concentration in mg ml–1)]. Graphs were plotted as relative EA normalized with respect to control. Assessment of Complex V activity was conducted using ATP synthase EA Kit (Abcam, Cat.: ab109714) as per Manufacture’s protocol. Relative EA was calculated as: (OD1 – OD2)/time. Graphs were plotted as relative EA normalized with respect to control.

### 2.6. Dopamine- and acetylcholine-dependent *C. elegans* Behavioral Assay

DA-dependent behavior (as an indirect measure of DA levels that have been validated pharmacologically), was quantified using 1-nonaol assay as described (Sammi et al. 2019). The relationship between DA levels and repulsion time has been repeatedly shown in the literature. 1-nonanol assay relies on the fact that nematodes exhibit repulsive behavior towards certain odorant chemicals which is inversely proportional to DA levels (Baidya et al. 2014; Kimura et al. 2010; Sammi et al. 2018b),. Conventionally, worms with lower levels of DA exhibit a longer repulsion time (Sammi et al. 2018a; Smita et al. 2017). Briefly, treated worms were washed 3 times with M9 buffer. Worms were placed on NGM plates. The poking lash dipped in 1-nonanol (Acros Organic, NJ, USA, AC157471000) was placed close to the head of the worms. Any worm prodded accidentally was disregarded. Time taken for the worms to show repulsive behavior was counted using a stopwatch.

Determination of acetylcholine-based function was conducted as described (Mahoney et al. 2006). The aldicarb assay relies on acetylcholinesterase AchE inhibition mediated paralysis in worms, where percentage of worms paralyzed is proportional to ACh levels (Mahoney et al. 2006; Sammi et al. 2019) Briefly, treated worms were washed three times and transferred to NGM-aldicarb plates (0.5 mM aldicarb). The percentage of worms paralyzed at a time point is proportional to ACh levels. The percentage of worms paralyzed was calculated when approximately 50% of worms were paralyzed in the control group.

### 2.7 RNA Isolation, cDNA Synthesis, and Quantitative Real Time PCR

Total RNA was extracted from worms exposed to CPF for 72 hrs using RNAzol reagent (Molecular Research Centre Inc.) according to the manufacturer’s instructions. Briefly, worms were washed with M9 buffer and crushed in RNAzol, followed by incubation at room temperature for 10 minutes. This was followed by centrifugation at 13000 rpm for 10 minutes at 4°C. The supernatant was collected and mixed with 100μl of nuclease free water. The vials were vortexed for 15s and kept at room temperature for 15 minutes followed by centrifugation at 13000 rpm at 4°C. RNA was precipitated from the supernatant by adding equal amount of isopropanol. The vials were centrifuged at 13000 rpm for 10 minutes and the pellet containing RNA was washed twice using 75% chilled ethanol and centrifuged at 6000 rpm for 50 minutes at 4°C. The pellet was dried and dissolved in nuclease free water followed by quantification using nanodrop spectrophotometer. cDNA synthesis was done from 1 μg of total *C. elegans* RNA in a thermal cycler using RevertAid H minus First Strand cDNA synthesis Kit (Thermo Scientific) according to manufacturer’s protocol. qRT-PCR studies were done using BioRad CFX 96. Differential expression was calculated by 2^-ΔΔCT^ method (Livak and Schmittgen 2001). *Gpd-1* was used as internal control and used to calculate fold change in mRNA expression. Primers were procured from Integrated DNA technologies, with details as shown in table 1 (orientation of oligonucleotides: 5’ to 3’).

**Table 1:**
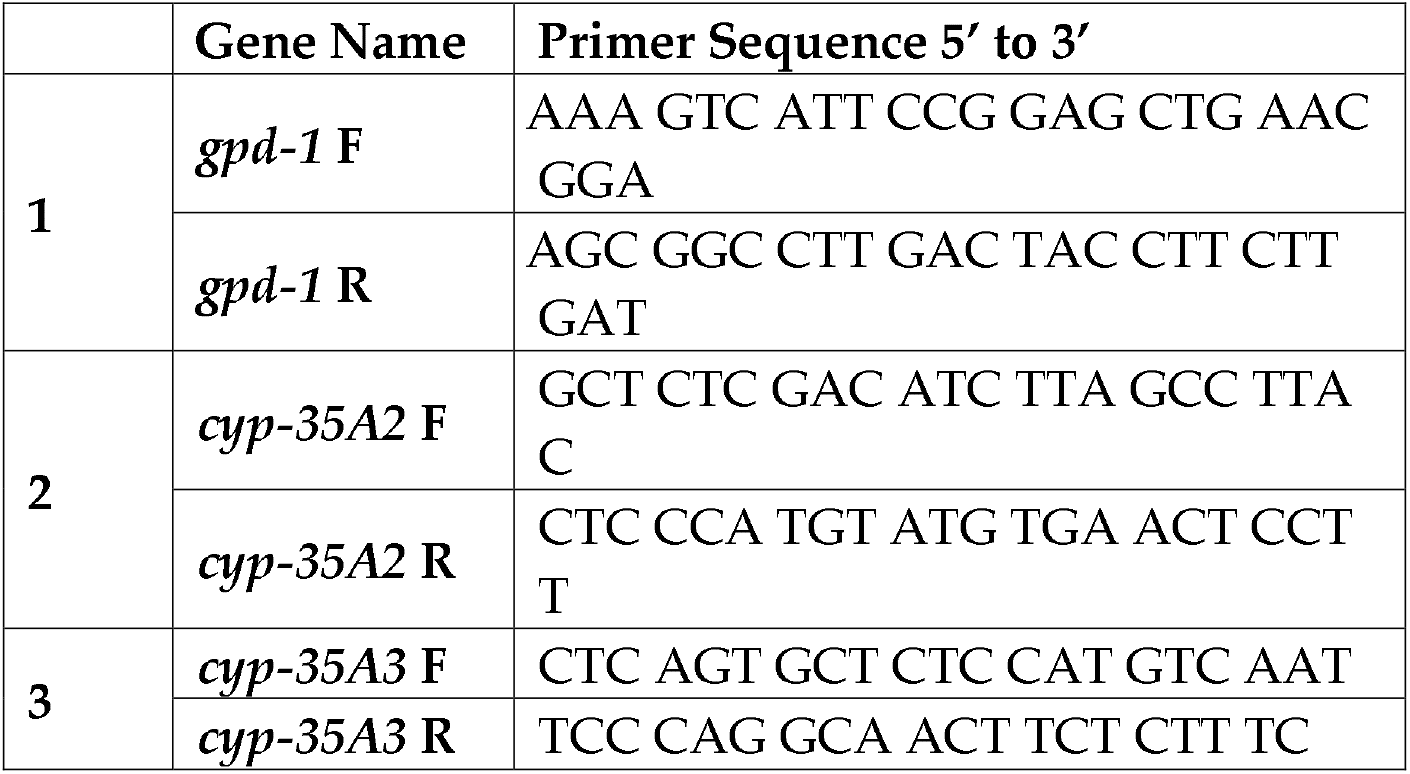
Sequence of the primers used.

### 2.8. Statistical Analysis of *C. elegans* endpoints and mitochondrial assays

Statistical analysis was performed using GraphPad PRISM, Version 7.02 (1992–2016 GraphPad Software, Inc., La Jolla, CA, USA). Data are expressed as mean ± S.E.M. Analysis of variance (ANOVA), followed by Dunnett’s (where comparison was only to control) and Sidak’s (in case of two-way ANOVA) *post hoc* tests were utilized. For all experiments, *p* < .05 was deemed statistically significant.

### 2.9. Examination Set of U.S. EPA ToxCast Assays

We examined all ToxCast assays tested on CPF, malathion, diazinon, and their oxon metabolites (i.e. CPF oxon, malaoxon, and diazoxon). We downloaded the assay information and data at the U.S. EPA CompTox Chemicals Dashboard (https://comptox.epa.gov/dashboard/) on March 9^th^, 2022. The assay information and data included a.) the name, target family, and gene annotation of each ToxCast assay and b.) the half-maximal activity concentrations (AC_50_s) of all 6 chemicals. We included the ToxCast assays from the target families of “nuclear receptor”, “cyp”, “oxidase”, “oxidoreductase”, “transferase”, and “transporter” to capture the bioactivity specific to gene targets in nuclear receptor signaling and fatty acid/steroid metabolism in our analysis. We also included the Tox21 HepG2 cell mitochondrial assays (Sakamuru *et al*., 2012), the *in vitro* neurodevelopment assays (Kosnik *et al*., 2020), and the microelectrode array assays (Harrill *et al*., 2018) to capture the cell-level events related to neurotoxicity. The calculation of AC_50_s was described in details by Judson et al (Judson et al. 2016). We excluded the ToxCast assays that did not have an AC_50_ value (i.e. negative results for all six chemicals tested).

### 2.10. Toxicity Score, Cluster Analysis, and ToxPi Visualization

The AC_50_ values (in μM) of each chemical was transformed to a toxicity score (*T*) as a measurement of toxic effects *in vitro*, where *T* = −log_10_ (AC_50_ (in μM) / 1,000). The toxicity score for an AC_50_ of 1,000 μM was defined as zero, since any higher concentration *in vitro* would no longer be physiologically relevant. Two-way hierarchical cluster analysis was conducted with the toxicity scores of all ToxCast assays in the examination set using Ward’s (1963) clustering method in Rstudio version 1.4.1717. The toxicity score of each ToxCast gene target was visualized using the Toxicological Priority Index (ToxPi; Marvel *et al*., 2018). Each target was annotated with a functional category according to the NIH/NIAID (National Institutes of Health/National Institute of Allergy and Infectious Diseases) Database for Annotation, Visualization and Integrated Discovery (DAVID) Bioinformatics Resources (2021 Update; Sherman *et al*., 2022), using terms from significant (*p* < 0.05) functional annotation clusters generally based on pathways and biological processes. The central angle of each slice in ToxPi indicated the number of ToxCast gene targets in each functional category. The length of each slice indicated the highest toxicity score in each functional category, which reflected the magnitude of effect regardless of its directionality. The cell-level events related to neurotoxicity were visualized in a separate ToxPi, where the central angle of each slice indicated the number of ToxCast assays dedicated to each event.

## 3. Results

### 3.1. CPF leads to dopaminergic cell loss in *C. elegans*

In view of the emerging evidence about DA toxicity, we tested different doses of CPF for DA cell loss in *C. elegans*. Nematodes exhibited a dose-dependent loss in DA neurons with variation in susceptibility amongst DA subpopulations (***Figure 1A***). Besides DA cell loss, worms also exhibited developmental delay (Supplementary figure S1, Table 1).

**Figure 1:**
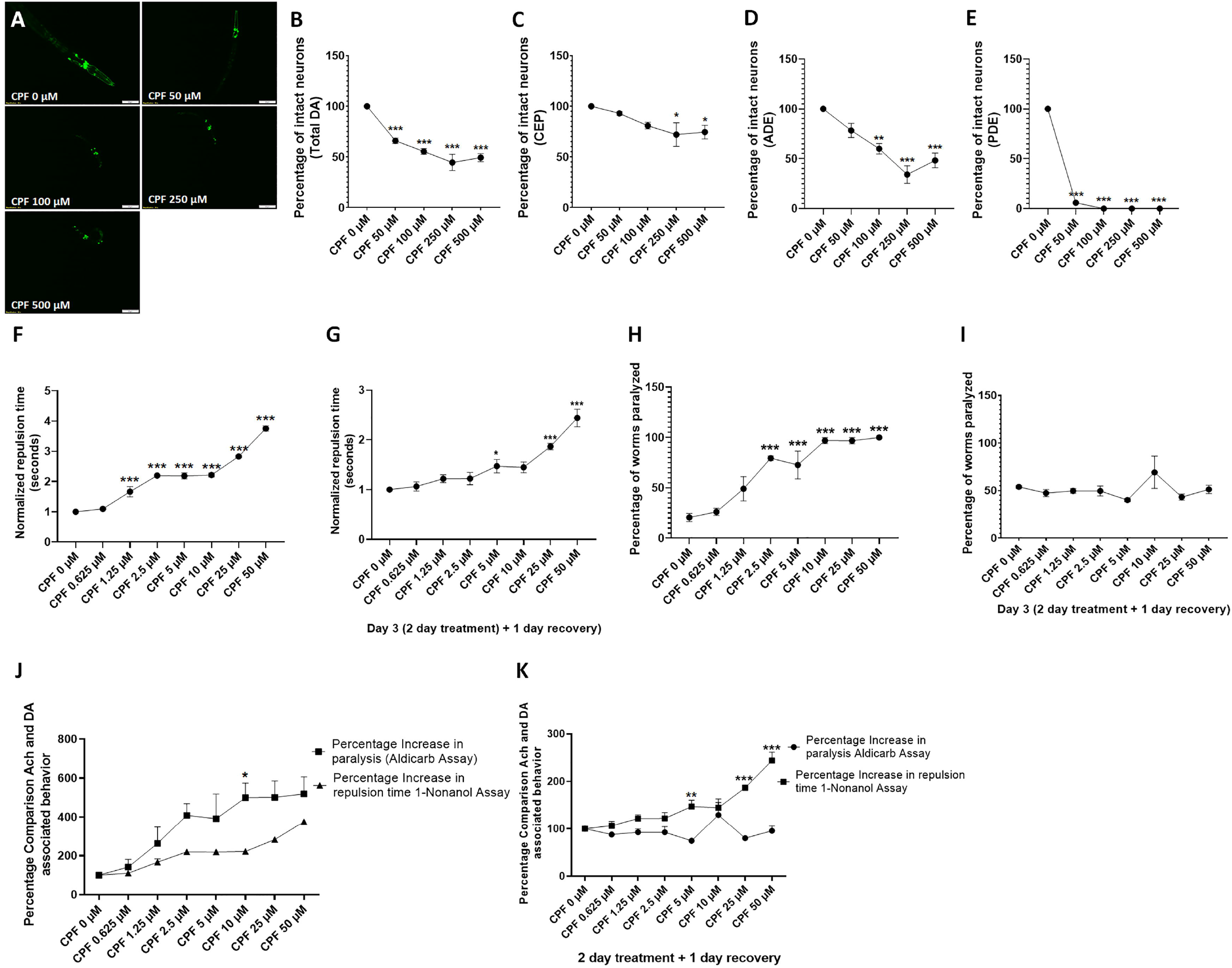
CPF confers DA neurotoxicity and alters dopamine and ACh associated behavior. Treatment of worms with CPF (exposure level range: 50 to 500 μM for 72 hrs) resulted in distinct morphological alterations such as axon breaks or loss of dendrites, swelling, and loss of soma, which represent neuronal damage in worms (***A***). The percentage of intact neurons calculated with respect to total neurons (***B***), CEP neurons (***C***), ADE neurons (***D***) and PDE neurons (***E***). A two day treatment with CPF produced increased repulsion time in response to 1-nonanol, indicative of lowered DA levels at concentrations 1.25 μM and above (***F***). Worms exhibited recovery from the effects of CPF when treated for two days and allowed to recover (without CPF) for one day. However higher concentrations retained the effects and exhibit significant increase in repulsion time at 5, 25 and 50 μM (***G***). CPF treatment produced a dose dependent increase in Aldicarb induced paralysis, indicative of higher synaptic ACh concentration at exposure levels 2.5 μM and above (***H***). Worms exhibited complete recovery from the effects of CPF when treated for two days and allowed to recover (without CPF) for one day (***I***). ACh and DA associated behavior without recovery (***J***) and with recovery (***K***) show a lack of reversivbility with DA neurotoxicity, whereas ACh deficits were reversible. Data are presented as mean ± S.E.M. The percentage of intact neurons was calculated by counting the total number of neurons in each worm for 20 worms per experimental group. Data were analyzed using one-way ANOVA followed by Dunnett’s post hoc test (***B–I***) and Sidak’s (***J–K***) post hoc tests. *p<.05, **p<.005, and ***p<.001 (n =3). Scale bar represents 50 μm

A significant reduction in percentage of intact neurons (PIN) was observed at doses 50 μM (66.042 ± 2.610, *p* =0.0009), 100 μM (55.417 ± 2.917, *p* =0.0001), 250 μM (44.583 ± 7.941, *p* <0.0001), and 500 μM (49.375 ± 4.018, *p* <0.0001) with respect to control (**Figure 1B**). CEP neurons showed least vulnerability with decrease in PIN at concentrations 250 μM (72.083 ± 11.555, *p* =0.0310), and 500 μM (74.583 ± 6.706, *p* =0.0494) as compared to control (***Figure 1C***). ADE neurons exhibited decrease in PIN at doses 100 μM (60.000 ± 5.204, *p* =0.0048), 250 μM (34.167 ± 8.819, *p* =0.0001), and 500 μM (48.333 ± 7.407, *p* =0.0008) in comparison to control (***Figure 1D***). PDE neurons found to be most vulnerable, exhibited reduction in PIN at 50 μM (5.833 ± 0.833, *p* <0.0001), 100 μM (0.000 ± 0.000, *p* <0.0001), 250 μM (0.000 ± 0.000, *p* <0.0001), and 500 μM (0.000 ± 0.000, *p* <0.0001) in comparison to control (***Figure 1E***). Taken together, our results confirmed DA degeneration in response to CPF.

### 3.2. Chlorpyrifos affects dopamine and acetylcholine dependent behavior differently

CPF acts as an AchE inhibitor (Costa et al. 2008b). In light of the observed DA neurotoxicity and reported cholinergic toxicity, we tested the effect of CPF on DA and ACh-associated behavior using 1-nonanol and aldicarb assay respectively. Two different treatment approaches were adopted: 1) exposing nematodes to CPF for 48 hrs, and 2) exposing nematodes to CPF for 48 hrs. followed by recovery for 24 hrs. (Washout experiments). Low doses of toxicants were employed to limit the bias resulting from toxicity mediated aberrations in motility. CPF treatment led to a significant increase in repulsion time at doses 1.25 μM (1.666 ± 0.169, *p* =0.0002), 2.5 μM (2.199 ± 0.050, *p* <0.0001), 5 μM (2.190 ± 0.099, *p* <0.0001), 10 μM (2.218 ± 0.063, *p* <0.0001), 25 μM (2.830 ± 0.040, *p* <0.0001) and 50 μM (3.753 ± 0.086, *p* <0.0001) in comparison to control, indicating a reduction of DA levels (***Figure 1F***). As expected, washout experiments showed recovery of DA deficit at lower doses, while higher doses, 5 μM (1.471 ± 0.135, *p* =0.0385), 25 μM (1.866 ± 0.063, *p* =0.0003) and 50 μM (2.439 ± 0.175, *p* <0.0001) retained the effect (***Figure 1G***).

Similarly, CPF treated worms exhibited increased paralysis, indicating increased ACh levels. Worms treated with CPF showed an increase in percentage of worms paralyzed at doses 2.5 μM (1.471 ± 0.135, *p* =0.0001), 5 μM (1.866 ± 0.063, *p* =0.0005), 10 μM (2.439 ± 0.175, *p* <0.0001), 25 μM (1.866 ± 0.063, *p* <0.0001) and 50 μM (1.866 ± 0.063, *p* <0.0001) in comparison to that of control (20.614 ± 4.021) (***Figure 1H***). In contrast to the washout experiments for 1-nonanol assay, we observed complete recovery of ACh-associated behavior (***Figure 1I***). This implied that the effect on DA-associated behavior persists even after withdrawal of the CPF. A comparison between the effect on DA and ACh associated behavior was made. For 48 hrs. studies, we observed a significant increase in percentage of paralyzed for ACh associated behavior at 10 μM (499.495 ± 74.875, *p* <0.0101) in comparison to a percentage increase in repulsion time 10 μM (221.835 ± 6.341) (***Figure 1J***). In case of washout experiments, we observed a significant increase of percentage increase in repulsion time at doses 5 μM (147.118 ± 13.531, *p* <0.0018), 25 μM (186.591 ± 6.284, *p* <0.0001), and 50 μM (243.910 ± 17.531, *p* <0.0001) as compared to the percentage increase in paralysis at doses 5 μM (74.430 ± 3.850), 25 μM (80.048 ± 4.898), and 50 μM (95.572 ± 10.227) respectively (***Figure 1K***). Collectively, our results demonstrated that CPF alters DA and ACh associated behavior. While alteration in ACh associated behavior is more than DA behavior, the later persists even after CPF is no longer present.

### 3.3. CPF treatment exhibits loss of mitochondrial content

Mitochondrial dysfunction is a critical primary PD mechanism (Park et al. 2018). PD-related toxicants such as rotenone directly target mitochondrial function, where dopaminergic neurons are especially sensitive (Chernivec et al. 2018; Greenamyre et al. 2003). Hence, we evaluated the effect of CPF on mitochondrial content in C. *elegans* using the reduced form of mitotracker red (MitoTracker Red CM-H2Xros), which stains viable mitochondria depending on membrane potential in live cells, (Invitrogen 2008). Worms treated with CPF and stained with Mito Tracker red exhibited decreased content of viable mitochondria (normalized with respect to control) in a dose-dependent fashion at the concentration 50 μM (0.539 ± 0.041, *p* =0.0002), 100 μM (0.103 ± 0.047, *p* <0.0001), 250 μM (0.108 ± 0.092, *p* <0.0001), and 500 μM (0.016 ± 0.001, *p* <0.0001) with respect to control (***Figure 2 A, B***). These results implied deleterious effects of CPF on mitochondria.

**Figure 2:**
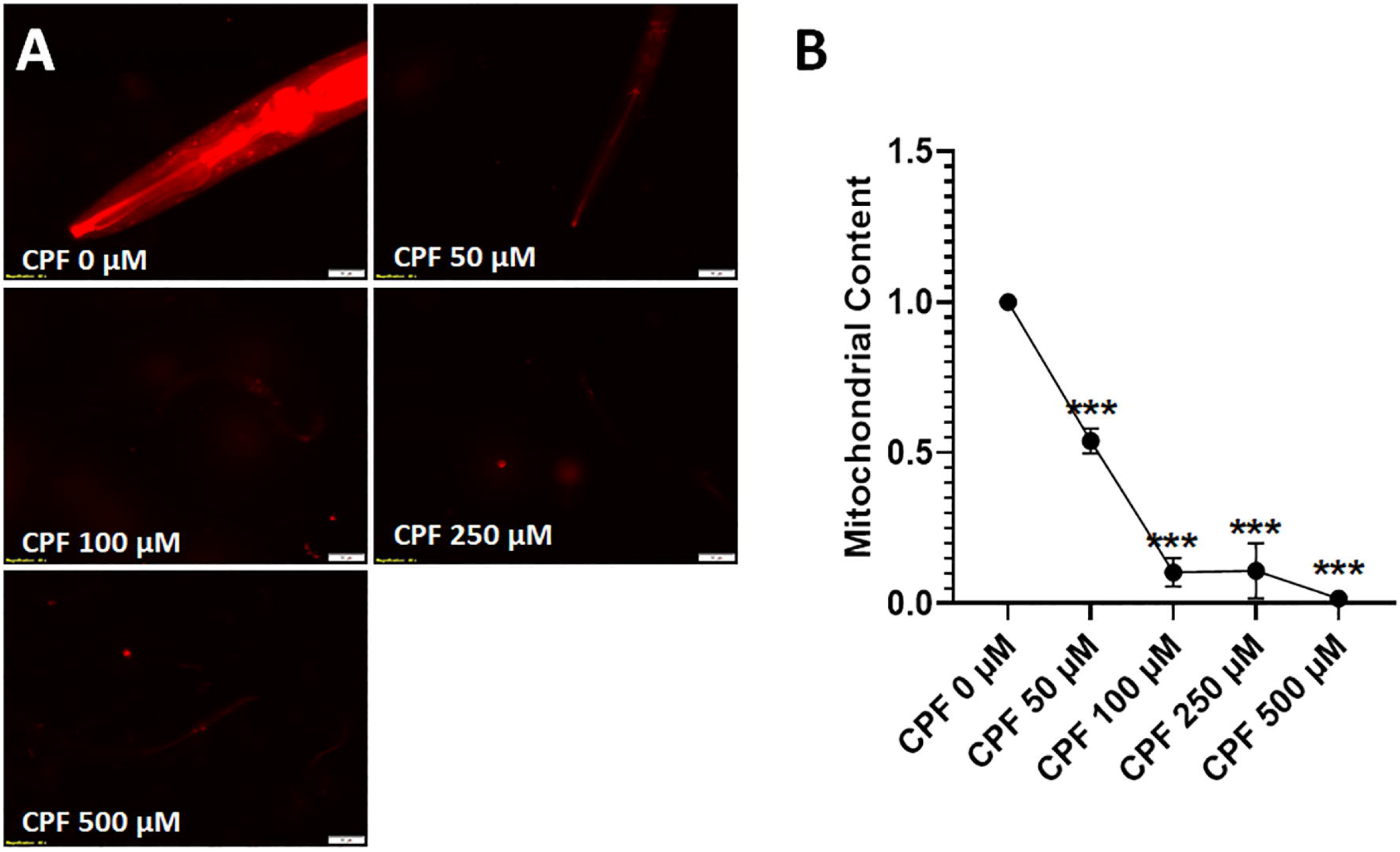
CPF damages mitochondria. CPF treated worms exhibited reduced mitochondrial content in an exposure dependent manner (***A***). Significant decrease in mitochondrial content was observed at doses 50 μM and above (***B***). Mitochondrial content was semi-quantitatively calculated through Image J for 20 worms per experimental group. Data were analyzed using one-way ANOVA followed by Dunnett’s post hoc test. ***p<.001 (n =3). Scale bar represents 20 μm (A). Data were analyzed using one-way ANOVA followed by Dunnett’s post hoc test. ***p<.001 (n =3).

### 3.4. CPF metabolite, CPF oxon, is more toxic than CPF

CPF oxon is the widely believed to be a primary mediator of CPF neurotoxicity (Supreeth and Raju 2017; Wang et al. 2010). Indeed, this was found to be true with respect potency to dopaminergic neurotoxicity. We tested effects of CPF vs CPF-Oxon was tested for effect on DA cell loss and DA associated behavior. A significant difference in PIN for total neurons was observed in case worms treated with CPF-Oxon 12.5 μM (51.042 ± 2.456, *p* < *0.0001*) in comparison to CPF 12.5 μM (97.500 ± 0.955); CPF-Oxon 25 μM (42.500 ± 6.683, *p* < *0.0001*) in comparison to CPF 25 μM (82.917 ± 2.894); CPF-Oxon 50 μM (20.417 ± 5.640, *p* < 0.0001) in comparison to CPF 50 μM (81.042 ± 6.706); CPF-Oxon 100 μM (17.708 ± 0.833, *p* < 0.0001) in comparison to CPF 100 μM (61.667 ± 2.456); CPF-Oxon 250 μM (8.750 ± 5.316, *p* < *0.0001*) in comparison to CPF 250 μM (53.333 ± 8.007); CPF-Oxon 500 μM (0.000 ± 0.000, *p* < *0.0001*) in comparison to CPF 500 μM (41.042 ± 3.858) (***Figure 3A, B***).

**Figure 3:**
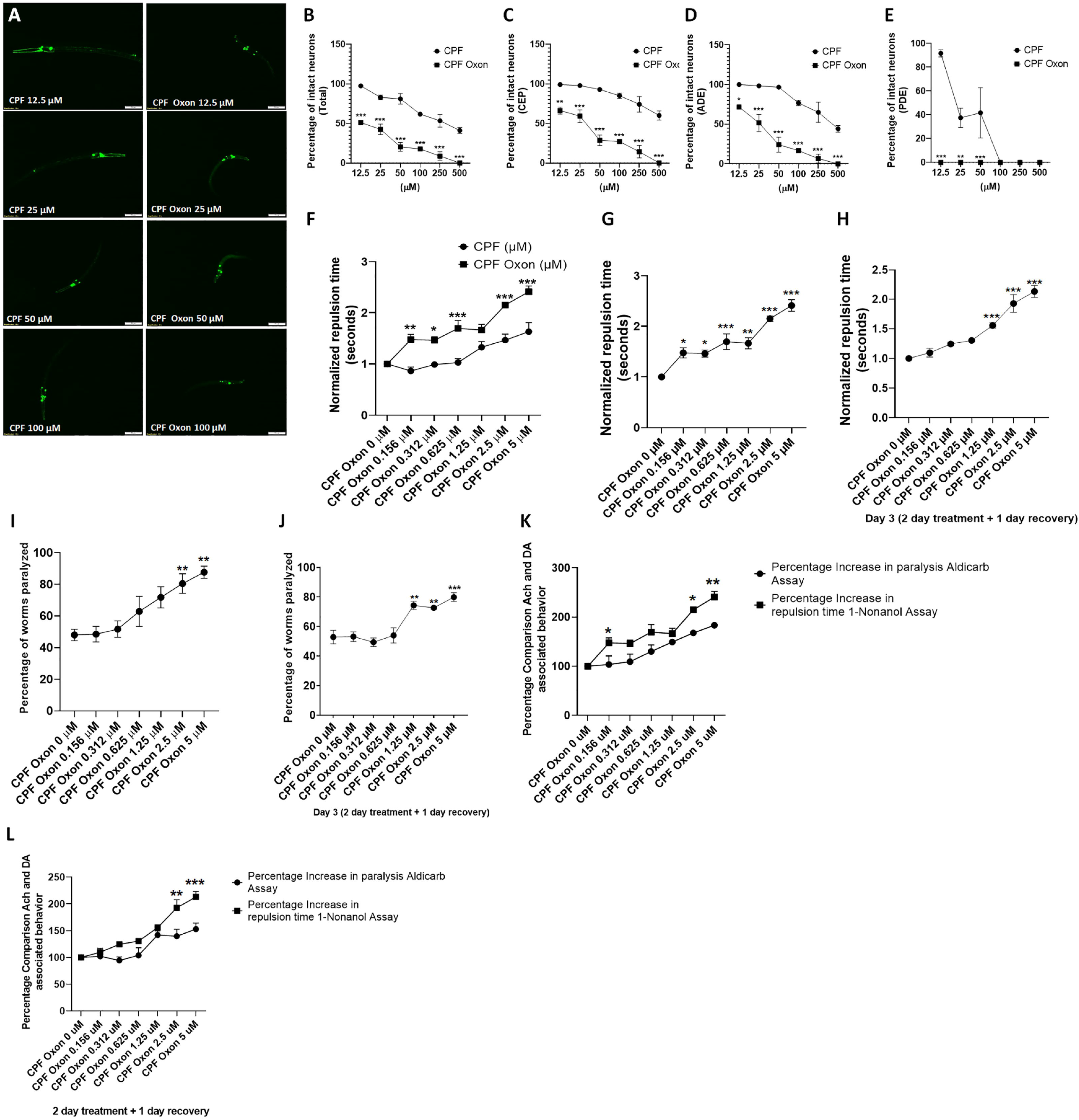
CPF oxon is highly potent to DA neurons and associated behavior. Representative images? Treatment of worms with CPF oxon (exposure level range: 12.5 to 500 μM for 48 hrs) resulted in distinct morphological changes such as axon breaks or loss of dendrites, swelling, and loss of soma, characteristic of neuronal damage in worms (***A***). Quantification of effect of CPF and CPF oxon on the percentage of intact neurons calculated with respect to total neurons, (showing high oxon potency in comparison to CPF) (***B***) in total DA neurons and specific subpopulations: CEP neurons (***C***), ADE neurons (***D***) and PDE neurons (***E***). Data are presented as mean ± S.E.M. The percentage of intact neurons was calculated by counting the total number of neurons in each worm for 20 worms per experimental group. Comparison of effect of CPF oxon Vs CPF treatment on DA dependent behavior (***F***). A two day treatment with CPF oxon exhibited increase in repulsion time in response to 1-nonanol, indicative of lowered DA levels at concentrations 0.156 μM and above (***G***). Worms exhibited slight recovery from the effects of CPF oxon when treated for two days and allowed to recover (without CPF oxon) for one day. However, higher concentrations retained the effects and exhibit significant increase in repulsion time at 1.25 to 5 μM (***H***). CPF oxon treatment showed a dose dependent increase in Aldicarb induced paralysis, indicative of higher synaptic ACh concentration at exposure levels 2.5 μM and above (I). Worms exhibited slight recovery form the effects of CPF oxon when treated for two days and allowed to recover (without CPF oxon) for one day (***J***). Comparison of ACh and DA associated behavior without recovery (***K***) and with recovery (***L***). Data analyzed using one-way ANOVA followed by Sidak’s (***B–F, K–L***) and Dunnett’s (***G – J***) post hoc test. *p<.05, **p<.005, and ***p<.001 (n =3). Scale bar represents 50 μm (A).

A significant difference in PIN for CEP neurons was observed in case worms treated with CPF-Oxon 12.5 μM (66.250 ± 4.390, *p* = *0.0012*) in comparison to CPF 12.5 μM (99.167 ± 0.417); CPF-Oxon 25 μM (59.167 ± 7.982, *p* = *0.0002*) in comparison to CPF 25 μM (97.917 ± 1.102); CPF-Oxon 50 μM (28.750 ± 6.614, *p* < *0.0001*) in comparison to CPF 50 μM (92.917 ± 2.083); CPF-Oxon 100 μM (27.083 ± 0.833, *p* < *0.0001*) in comparison to CPF 100 μM (85.000 ± 3.307); CPF-Oxon 250 μM (14.167 ± 7.949, *p* < *0.0001*) in comparison to CPF 250 μM (74.167 ± 9.851); CPF-Oxon 500 μM (0.000 ± 0.000, *p* < *0.0001*) in comparison to CPF 500 μM (60.000 ± 5.637) (***Figure 3A, C***).

A significant difference in PIN for ADE neurons was observed in case worms treated with CPF-Oxon 12.5 μM (71.667 ± 1.667, *p* = 0.0168) in comparison to CPF 12.5 μM (100.000 ± 0.000); CPF-Oxon 25 μM (51.667 ± 10.833, *p* < *0.0001*) in comparison to CPF 25 μM (98.333 ± 1.667); CPF-Oxon 50 μM (24.167 ± 9.391, *p* < *0.0001*) in comparison to CPF 50 μM (96.667 ± 2.205); CPF-Oxon 100 μM (16.667 ± 1.667, *p* < *0.0001*) in comparison to CPF 100 μM (76.667 ± 3.333); CPF-Oxon 250 μM (6.667 ± 5.465, *p* < *0.0001*) in comparison to CPF 250 μM (65.000 ± 12.583); CPF-Oxon 500 μM (0.000 ± 0.000, *p* = 0.0002) in comparison to CPF 500 μM (44.167 ± 4.167) (***Figure 3A, D***).

A significant difference in PIN for PDE neurons was observed in case worms treated with CPF-Oxon 12.5 μM (0.000 ± 0.000, *p* < *0.0001*) in comparison to CPF 12.5 μM (97.500 ± 0.955); CPF-Oxon 25 μM (0.000 ± 0.000, *p* = *0.0029*) in comparison to CPF 25 μM (82.917 ± 2.894); CPF-Oxon 50 μM (0.000 ± 0.000, *p* = *0.0009*) in comparison to CPF 50 μM (81.042 ± 6.706); (***Figure 3A, E***).

### 3.5. CPF Oxon alters dopamine and acetylcholine associated behavior

Given that CPF Oxon is more toxic than the parent compound, CPF, we compared the effect of CPF-Oxon with CPF on DA associated behavior and also elucidated the comparison between DA and ACh associated behavior.

Due to the high toxicity of CPF-Oxon, a lower dose range was selected (0.156 μM to 5 μM). CPF-Oxon exhibited increased repulsion time, indicative of lower DA levels. In comparison to CPF 0.156 μM (0.863 ± 0.074), CPF 0.312 μM (0.989 ± 0.038), CPF 0.625 μM (1.028 ± 0.076), CPF 2.5 μM (1.465 ± 0.116), and CPF 5 μM (1.630 ± 0.176) a significant increase in normalized repulsion time was observed in case of CPF-Oxon 0.156 μM (1.475 ± 0.101, *p* = 0.0010), CPF-Oxon 0.312 μM (1.461 ± 0.070, *p* = 0.0145), CPF-Oxon 0.625 μM (1.694 ± 0.153, *p* = 0.0004), CPF-Oxon 2.5 μM (2.149 ± 0.052, *p* = 0.0002), and CPF-Oxon 5 μM (2.407 ± 0.113, *p* < 0.0001) respectively (***Figure 3F***).

CPF oxon treatment led to a significant increase in normalized repulsion time at CPF oxon 0.156 μM (1.475 ± 0.101, *p* = 0.0182), CPF oxon 0.312 μM (1.461 ± 0.070, *p* = 0.0221), CPF oxon 0.625 μM (1.694 ± 0.153, *p* = 0.0009), CPF oxon 1.25 μM (1.663 ± 0.110, *p =* 0.0014), CPF oxon 2.5 μM (2.149 ± 0.052, *p* < 0.0001), and CPF oxon 5 μM (2.407 ± 0.113, *p* < 0.0001) as compared to that of CPF oxon 0 μM (1.000 ± 0.000) (***Figure 3G***). Washout experiments, where worms were allowed to recover for a 24 hrs post 48 hrs treatments showed slight recovery in worms. In comparison to the normalized repulsion time in control (CPF oxon 0 μM) (1.000 ± 0.000), a significant increase in repulsion time was seen in worm treated with CPF oxon 1.25 μM (1.557 ± 0.043, *p* = 0.0008), CPF oxon 2.5 μM (1.928 ± 0.149, *p* < 0.0001), and CPF oxon 5 μM (2.131 ± 0.100, *p* < 0.0001) (***Figure 3H***). This implied the CPF oxon led to decline in DA levels at lower doses, and the effect was persistent even after worms were allowed to recover for 24 hrs.

Similarly, we also observed an increase in aldicarb induced paralysis, which relates to enhanced accumulation of ACh. In comparison to CPF oxon 0 μM (48.004 ± 3.625), a significant increase in percentage of worms paralyzed was observed at CPF oxon 2.5 μM (80.440 ± 6.149, *p* = 0.0088), and CPF oxon 5 μM (87.646 ± 3.813, *p* = 0.0018) (***Figure 3I***). In the washout experiments, where the worms were incubated in absence of CPF oxon for 24 hrs, we observed a significant increase in percentage of worms paralyzed at doses 1.25 μM (74.315 ± 2.784, *p* = 0.0030), 2.5 μM (72.685 ± 1.225, *p* = 0.0057), and 5 μM (79.894 ± 2.946, *p* = 0.0004) in comparison to that of 0 μM (52.862 ± 4.559) (***Figure 3J***).

In order to draw a comparison between the effect on DA and ACh associated behavior, we plotted the results from 1-nonanol and aldicarb assay against each other. For 48 hrs studies, we observed a significant decrease in percentage increase in paralysis for ACh associated behavior at CPF oxon 0.156 μM (103.623 ± 17.134, *p* = 0.0341) Vs percentage increase in repulsion time (147.507 ± 10.106), percentage increase in paralysis for ACh associated behavior CPF oxon 2.5 μM (167.753 ± 5.711, *p* = 0.0194) Vs percentage increase in repulsion time (214.895 ± 5.150), and percentage increase in paralysis for ACh associated behavior for CPF oxon 5 μM (183.431 ± 5.638, *p* = 0.0031) Vs percentage increase in repulsion time (240.736 ± 11.344) (***Figure 3K***).

In case of washout experiments, we observed a significant decrease in percentage increase in repulsion time at CPF oxon 2.5 μM (139.702 ± 12.922, *p* = 0.0018) Vs percentage increase in repulsion time (192.776 ± 14.947), CPF oxon 5 μM (152.944 ± 11.196, *p* = 0.0004) Vs percentage increase in repulsion time (213.097 ± 10.021) (***Figure 3L***). These results indicated that CPF-Oxon is far more toxic to DA neurons and DA associated behavior.

### 3.6. Effect of CPF and CPF oxon on mitochondrial complex activity

We performed assays for assessment of EA in mitochondria isolated from rat liver. Initial studies were conducted on three doses (25 to 75 μM). Detailed investigation using a broader dose range was conducted where significant effects were observed. Overall, complex I, III and IV were not found to be critical CPF targets (no significant inhibition up to 75 μM), while a significant effect of CPF was observed on complex II, II + III, and V (***Figure 4***). Notably mitochondrial complex II is a common component of both Electron transport chain and Tricarboxylic acid sycle, which also has a critical role in cell death and survival, while also acting as a source of reserve respiratory capacity (Pfleger et al. 2015). On the other hand mitochondrial complex III functions to oxidise the reduced Coenzyme and transfer the electrons to cytochrome C (Banerjee et al. 2021). Give the cooperation between Complex II, III and role of Coenzyme Q as a electron carrier, we also tested the Complex II + III activity, along with the effect of CoQ supplementation in enzyme activity.

**Figure 4:**
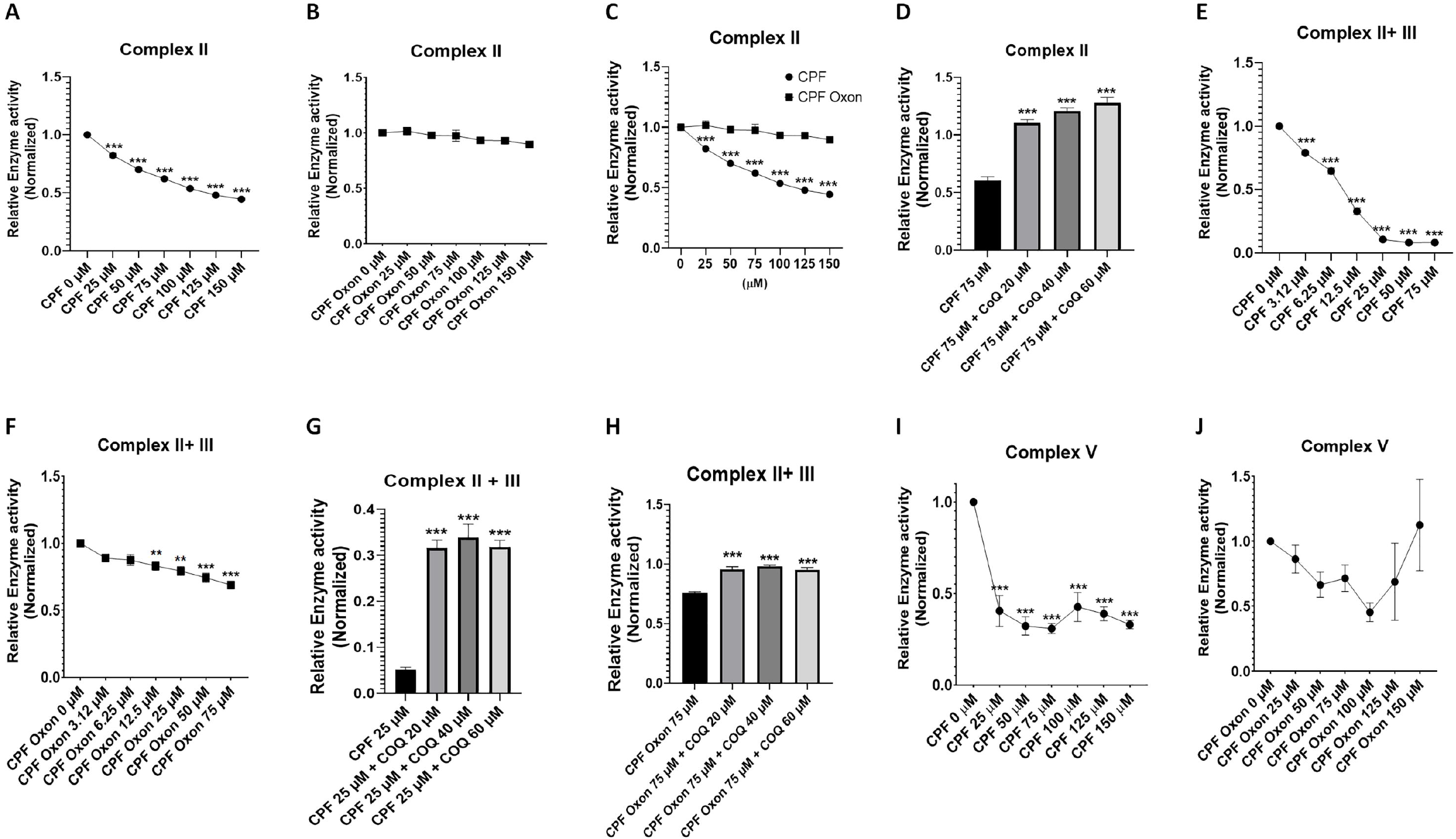
CPF targets specific complexes in the mitochondrial respiration chain. The effect of CPF and CPF oxon on mitochondrial respiration was assessed using mitochondria isolated from rat liver. **Complex II:** CPF treatment exhibited significant dose-dependent inhibition of Complex II activity (***A***), while CPF oxon lacked any effect (***B***). Direct comparisons of CPF and CPF oxon on Complex II activity show potency differences (***C***). Complex II activity inhibition was ameliorated by the electron acceptor CoQ (***D***); **Complex II + III:** CPF (***E***) and CPF oxon (***F***), both exhibited significant inhibiton of Complex II + III activity, with the effect more pronounced in case of CPF. As expected, CoQ treatment ameliorated inhibition of Complex II + III activity in both CPF (***G***) and CPF oxon (H); **Complex V:** CPF treatment (***I***) exhibited significant decrease in complex V activity, however there was absence of apparent dose dependent effect. CPF oxon lacked any significant effect on Complex V activity (***J***). Data analyzed using one-way ANOVA followed by Dunnett’s post hoc test. **p<.005, and ***p<.001 (n =3).

#### 3.6.1 CPF alters Complex II activity

CPF exhibited a dose dependent decrease in Complex II activity (***Figure 4A***) whereas CPF oxon was devoid of any effect on complex II activity (***Figure 4B***),. In comparison to relative EA in control (1.000 ± 0.000), a decrease in EA was observed at CPF 25 μM (0.822 ± 0.018, *p* < 0.0001), CPF 50 μM (0.700 ± 0.014, *p* < 0.0001), CPF 75 μM (0.620 ± 0.006, *p* < 0.0001), CPF 100 μM (0.536 ± 0.008, *p* < 0.0001), CPF 125 μM (0.479 ± 0.008, *p* < 0.0001), and CPF 150 μM (0.444 ±0.011, *p* < 0.0001) (Figure 4A). Comparison between the relative EA for CPF Vs CPF oxon shown in ***Figure 4C***.

#### 3.6.2 Coenzyme Q supplementation rescues Complex II activity in CPF treated mitochondria

Recent studies demonstrate that organophosphates mitigate Coenzyme (CoQ); CoQ supplementation exhibits a rescuing effect on enzyme activity (citrate synthase and Complex II+III) and cell viability (Turton et al. 2021). We observed significant amelioration of complex II activity at CPF 75 μM (0.603 ± 0.033) in comparison to CPF 75 μM co-treated with CoQ 20 μM (1.102 ± 0.033, *p* < 0.0001), CoQ 40 μM (1.207 ± 0.029, *p* < 0.0001), and CoQ 60 μM (1.277 ± 0.050, *p* < 0.0001) (***Figure 4D***). Thus, validating previous findings.

#### 3.6.3 CPF and CPF oxon inhibit Complex II + III activity

Next we tried to ascertain the effect of CPF and CPF oxon on EA for complex II + III. We observed a significant decrease in complex II + III EA in mitochondria treated with CPF 3.12 μM (0.789 ± 0.015, *p* < 0.0001), CPF 6.25 μM (0.646 ± 0.023, *p* < 0.0001), CPF 12.5 μM (0.328 ± 0.026, *p* < 0.0001), CPF 25 μM (0.106 ± 0.002, *p* < 0.0001), CPF 50 μM (0.082 ± 0.011, *p* < 0.0001), and CPF 75 μM (0.082 ± 0.005, *p* < 0.0001), in comparison to control (1.000 ± 0.000) (***Figure 4E***). Although to a lower extent, a decrease in complex II + III EA was also observed upon treatment with CPF oxon 12.5 μM (0.831 ± 0.032, *p* < 0.0001), CPF oxon 25 μM (0.794 ± 0.033, *p* < 0.0001), CPF oxon 50 μM (0.744 ± 0.035, *p* < 0.0001), and CPF oxon 75 μM (0.689 ± 0.025, *p* < 0.0001) (***Figure 4F***) as compared to control (1.000 ± 0.000). These results implied inhibitory effect of CPF and CPF oxon.

#### 3.6.4 CoQ supplementation rescues Complex II + III activity in CPF and CPF oxon treated mitochondria

CoQ supplementation alleviated the effect of CPF and CPF oxon. Compared to CPF 25 μM (0.015 ± 0.005), we observed a significant increase in complex II + III EA in CPF 25 μM + CoQ 20μM (0.316 ± 0.017, *p* < 0.0001), CPF 25 μM + CoQ 40μM (0.339 ± 0.029, *p* < 0.0001), and CPF 25 μM + CoQ 60μM (0.318 ± 0.014, *p* < 0.0001) (***Figure 4G***). Similarly, compared to CPF oxon 75 μM (0.755 ± 0.012), we observed a significant increase in complex II + III EA in mitochondria treated with CPF oxon 75 μM + CoQ 20μM (0.958 ± 0.021, *p* < 0.0001), CPF oxon 75 μM + CoQ 40μM (0.982 ± 0.009, *p* < 0.0001), and CPF oxon 75 μM + CoQ 60μM (0.952 ± 0.018, *p* < 0.0001) (***Figure 4H***).

#### 3.6.5 CPF alters complex V activity

CPF oxon lacked any effect on complex V EA (Figure 4J). CPF did alter complex V EA. In comparison to the control (1.000 ± 0.000), we observed a significant decrease in EA at doses, CPF 25 μM (0.405 ± 0.085, *p* < 0.0001), CPF 50 μM (0.322 ± 0.028, *p* < 0.0001), CPF 75 μM (0.308 ± 0.028, *p* < 0.0001), CPF 100 μM (0.426 ± 0.079, *p* < 0.0001), CPF 125 μM (0.389 ± 0.038, *p* < 0.0001), and CPF 150 μM (0.330 ± 0.023, *p* < 0.0001) (***Figure 4I***).

### 3.7. Effect of chlorpyrifos on expression of genes, *cyp-35A2*, and *cyp-35A3*

After determining the neurotoxic effects of CPF on DA neurons, we studied the effect of CPF on expression of genes related to organophosphate toxicity (*cyp-35A2* and *cyp35A3*) (*Roh and Choi 2011*; *Roh et al. 2014*). A significant dose dependent increase in mRNA expression of *cyp-35A2* was observed at CPF 3.12 μM (2.700 +0.224, *p* = 0.0004), and CPF 6.25 μM (3.689 ± 0.135, *p* < 0.0001) as compared to control (1.000 +0.000) (***Figure 5A***); A significant increase in mRNA expression of *cyp-35A3* was observed at CPF 6.25 μM (2.288 ± 0.353, *p* = 0.0127) as compared to control (1.000 +0.000) (***Figure 5B***). Lower doses of CPF were chosen to minimize the effects on gene expression incurred due to developmental delay.

**Figure 5:**
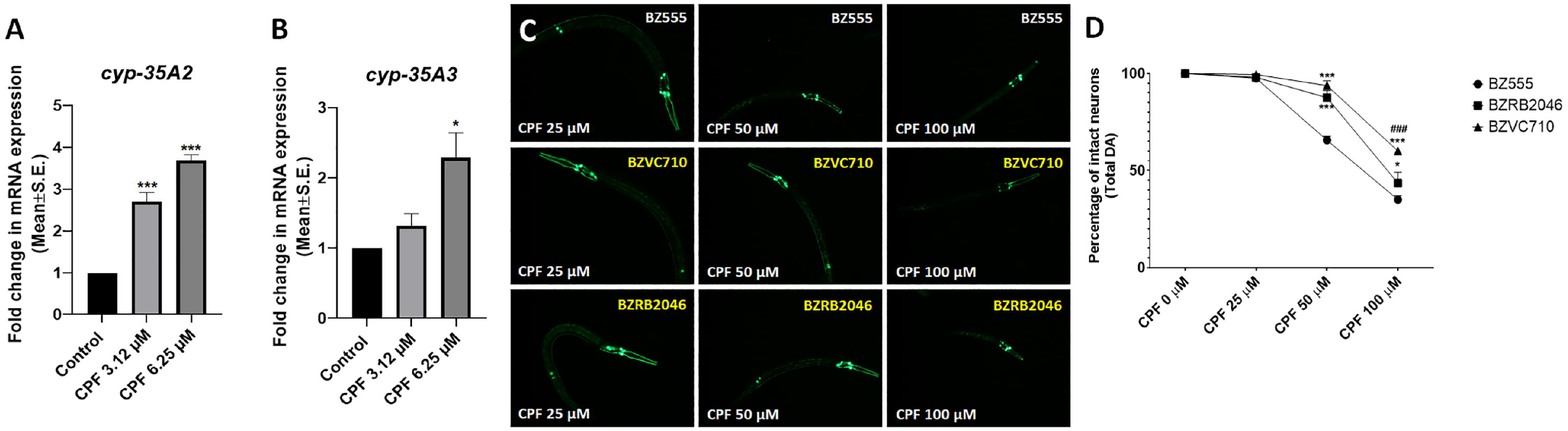
Cyp-35A2 and cyp-35A3 are critical modulators of CPF neurotoxicity. Worms treated at L1 stage with CPF exhibited increases in mRNA expression of *cyp-35A2* (***A***) and *cyp-35A3* (***B***). Mutants for *cyp-35A2* (BZVC710) and *cyp-35A3* (BZRB2046) exhibited neuroprotection (***C***) in comparison to BZ555 upon treatment with CPF (***D***). Data are presented as mean ± S.E.M. The percentage of intact neurons was calculated by counting the total number of neurons in each worm for 20 worms per experimental group. Data analyzed using one-way ANOVA followed by Dunnett’s (***A – B***) and Sidak’s (***D***) post hoc tests. *p<.05, and ***p<.001 (n =3). Scale bar represents 50 μm

### 3.8. Gene *cyp-35A2* and *cyp35A3* are critical for CPF neurotoxicity

After observing significant upregulation in expression of *cyp-35A2* and *cyp-35A3*, we studied the effect of mutation in these genes on DA neurotoxicity. Furthermore, *cyp-35a* family of genes, particularly *cyp-35A2* and *cyp-35A3* have been known to be upregulated in response to CPF exposure (Roh et al. 2014). Additionally *cyp-35A2* has been shown to be critical in conferring fenitrothioin (an organophosphate) toxicity (Roh and Choi 2011). Hence we crossed BZ555 with mutants for *cyp-35A2* (denoted as BZVC710) and *cyp-35A3* (denoted as BZRB2046) and studied the effect of these mutation on CPF mediated DA toxicity. The nematodes with mutation in *cyp-35A2* gene exhibited a significant increase in PIN at CPF 50 μM (93.750 ± 2.366, *p* < 0.0001), and 100 μM (60.000 ± 0.625, *p* < 0.0001) in comparison to BZ555 at CPF 50 μM (65.625 ± 2.009), and 100 μM (35.000 ± 2.009) (Figure 5C, D). The nematodes with mutation in *cyp-35A3* gene exhibited a significant increase in PIN at CPF 50 μM (87.500 ± 3.767, *p* < 0.0001), and 100 μM (43.542 ± 5.547, *p* = 0.0443) in comparison to BZ555 at CPF 50 μM (65.625 ± 2.009), and 100 μM (35.000 ± 2.009) (***Figure 5C, D***). These results indicated that the genes *cyp-35A2* and *cyp-35A3* are critical for CPF neurotoxicity.

### 3.9. Clustering of *in vitro* effects on ER, PXR, and PPAR gamma pathways of chlorpyrifos, malathion, diazinon, and their oxon metabolites with neurotoxicity

A total of 832 ToxCast assays were tested on CPF, malathion, diazinon, and their metabolites. 40.6 percent of the assays (338 in total) produced at least one positive result for all six chemicals tested. These included 84 ToxCast assays that capture the bioactivity specific to 52 gene targets in the following functional groups: 25 gene targets in nuclear receptor signaling, 13 gene targets in cytochrome P450 families, 8 gene targets in transporters, and six gene targets in other metabolic enzymes. These also included three Tox21 HepG2 cell mitochondrial assays, 16 *in vitro* neurodevelopment assays, and 26 microelectrode array assays. Overall, CPF, diazinon, and malathion shared a more similar *in vitro* activity profile in these 338 assays as compared to their oxon metabolites. CPF, CPF-oxon, diazinon, and malathion produced similar activity in ER, PXR, and PPAR gamma pathway as well as microelectrode arrays. Two-way hierarchical cluster analysis was conducted with the toxicity scores of those selected 129 ToxCast assays as shown in ***Figure 6***.

**Figure 6:**
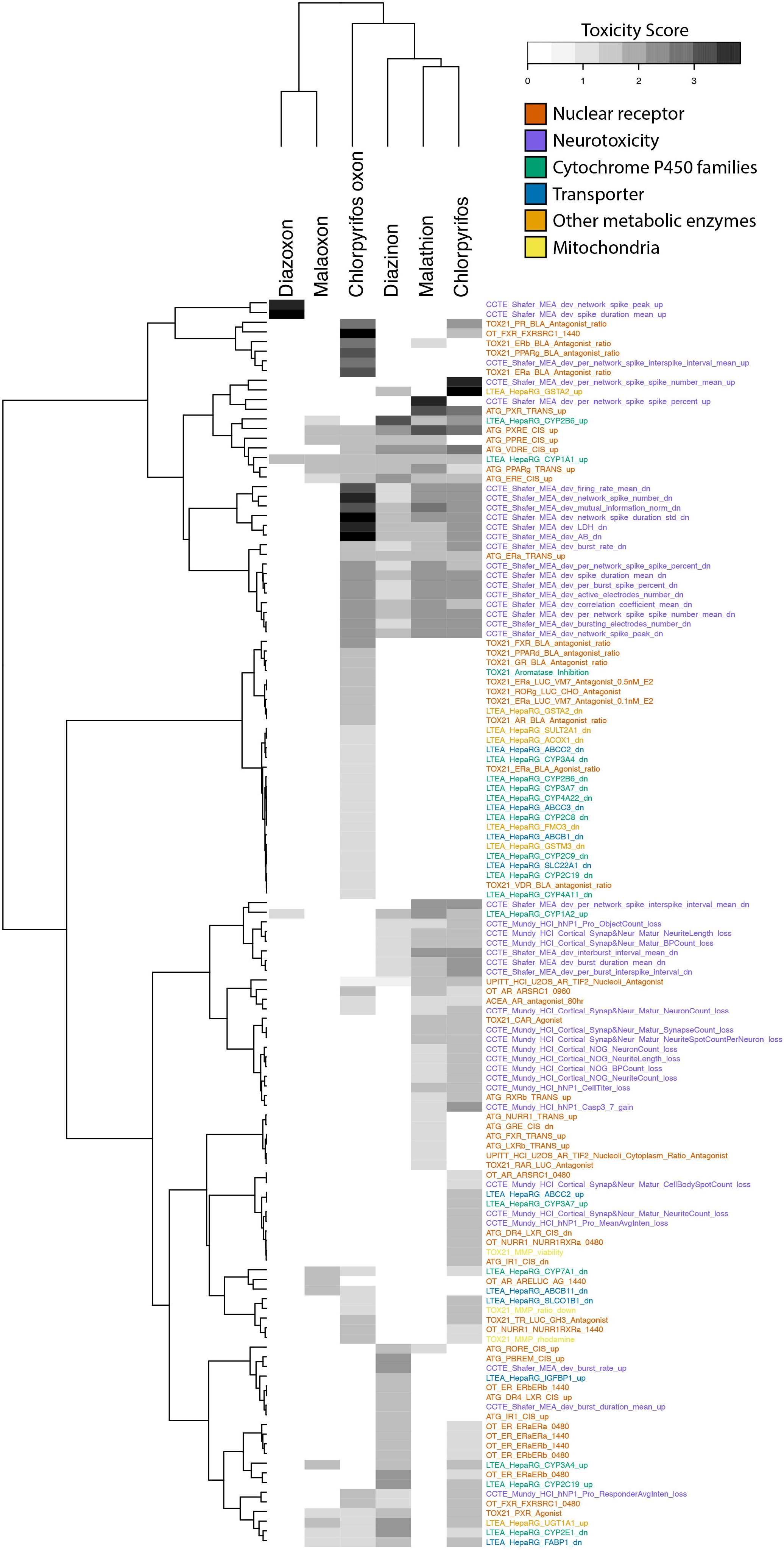
Ward clustering of the 129 selected ToxCast assays. 129 assays were selected from all ToxCast assays tested in all 6 OP/Oxon chemicals, which also had at least one positive assay result. 84 ToxCast assays were selected to capture the bioactivity specific to 52 gene targets, 42 assays related to neurotoxicity, and three assays were related to mitochondrial toxicity. The 52 gene targets were catergorized in the following functional groups: 25 gene targets in nuclear receptor signaling, 13 gene targets in cytochrome P450 families, 8 gene targets in transporters, and six gene targets in other metabolic enzymes. The assay names are colored according to these functional groups. Two-way hierarchical cluster analysis was conducted with the toxicity scores of those selected 129 ToxCast assays. Toxicity scores were calculated using −log_10_(AC50/1000).

### 3.10. Toxicity priority index for gene targets and cell-level events of CPF, malathion, diazinon, and their oxon metabolites

Selected assay gene targets were grouped using functional annotation with DAVID, and identified five nuclear receptor groups: steroid hormone receptors (23 assays), retinoic acid receptors (7 assays), xenobiotic metabolic receptors (5 assays), fatty acid metabolic receptors (4 assays), and other nuclear receptors (19 assays). Gene targets also included functional categories for CYP450 familes 1,2, or 3 (13 assays), other CYP450 families (4 assays), transporters (9 assays), and the other metabolic enzymes (7 assays). Cell level assay results were also grouped into functional categories including connectivity (14 assays), neurite outgrowth (6 assays), bursting (5 assays), cytotoxicity (5 assays), general electrical activity (5 assays), synaptogenesis (3 assays), proliferation (2 assays), apoptosis (1 assay), and mitochondrial (3) assays. The highest toxicity score within each functional group was chosen to visually represent the target toxicity as shown in ***Figure 7A and 7B***, each section sized by the number of assays that evaluate these functional groups.

**Figure 7:**
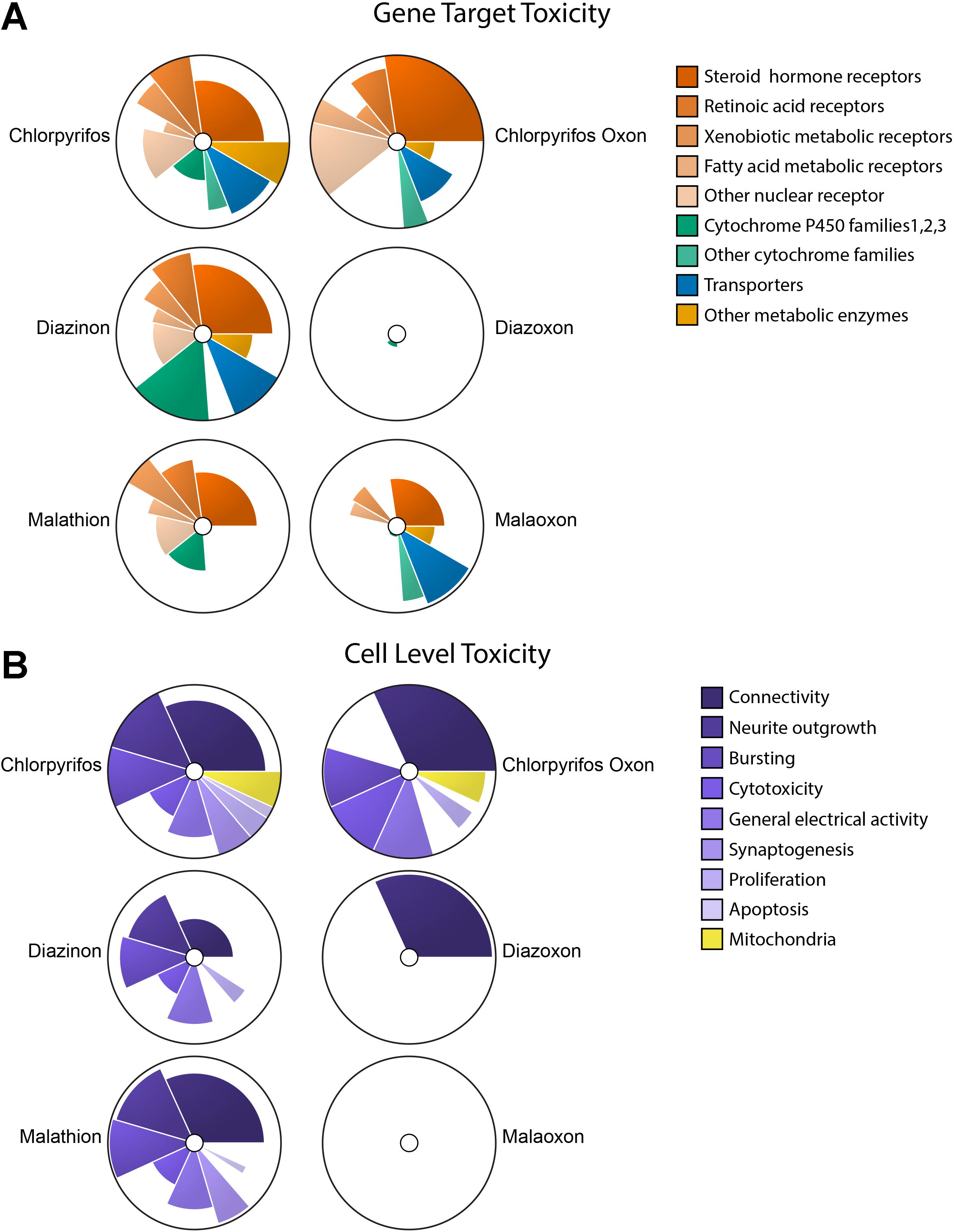
ToxPi visualization of highest toxicity scores for assay target groups. Toxicity scores for gene targets (***7A***) and cell level toxictiy (***7B***) for the 129 assays analyzed were split into two panels to vizualize toxicity in specific areas. Gene target slice angles were sized based on the number of gene targets each catergory contains, cell level toxicity slice angles were sized by the number of assays each catergory covered. Each radius is taken from the highest assay toxicity score to elimate the directionality of the assay results and elucidate main modes of actions more clearly than an average value calculation would. Each chart is colored by the main groupings and shaded by subcatergory.

**Figure 8:**
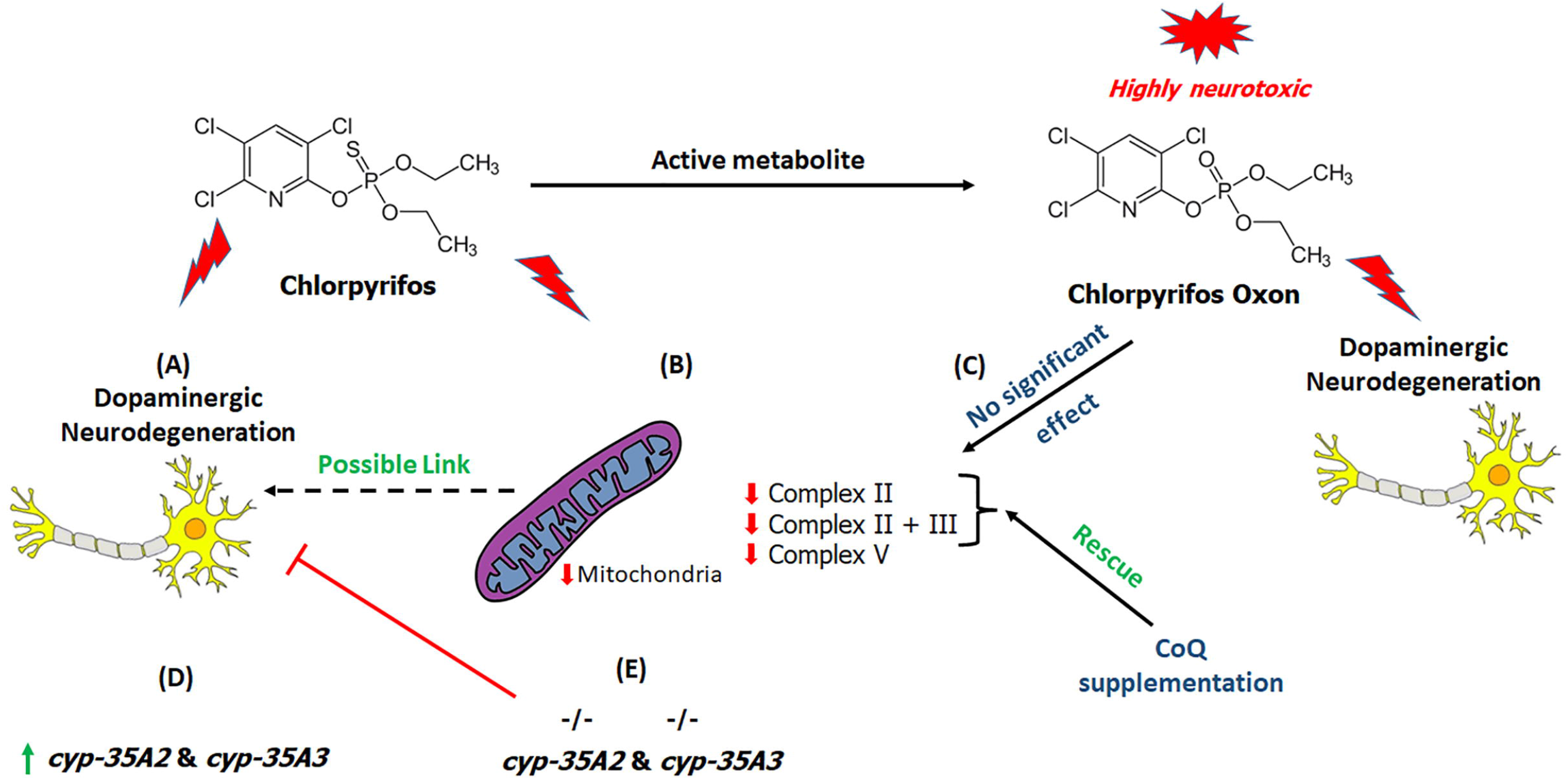
Proposed mechanism underlying chlorpyrifos DA neurotoxicity: Chlorpyriphos exerts dopaminergic neurotoxicity (***A***) through reduced mitochondrial content, inhibition of mitochondrial complexes II, II+III, and V. The effect on Complexes II, and II + III was reversed upon co treatment with CoQ (***B***). CPF is metabolized to CPF Oxon, which is highly neurotoxic yet lacks effect on mitochondrial complexes (***C***). CPF exposure induces upregulation of genes *cyp-35A2*, and *cyp-35A3* (***D***). Mutation in genes *cyp-35A2*, and *cyp-35A3* results in reduction of dopaminergic neurotoxicity (***E***). Overall, the study highlights the mechanisms pertaining to dopaminergic toxicity and mitochondrial toxicity, distinct from the mechanism knowns previously, particularly in the case of CPF metabolites and AchE inhibition.

## 4. Discussion

Chlorpyrifos is an extensively used, broad spectrum organophosphate pesticide, with effects emanating through inhibition of AchE (Maggio et al. 2021),(Burke et al. 2017). Upon entry into the body CPF is metabolized to CPF oxon by cytochrome P450 enzymes (Foxenberg et al. 2007; Tang et al. 2001). CPF oxon, a product of CPF desulfuration is highly neurotoxic (Tang et al. 2001). The ability to cross through blood brain barrier and target AchE renders CPF significantly neurotoxic to off-target organisms including mammals (Li and Ehrich 2013),(Levin et al. 2001). Thus, the majority of studies pertaining to CPF neurotoxicity focus on cholinergic neurons and AchE inhibition (Burke et al. 2017; Maggio et al. 2021). However, there is a significant knowledge gap pertaining to non-AchE neurotoxicity. Importantly, recent literature suggests potential risk to other neuron types such as DA neurons (Eddins et al. 2010; Singh et al. 2018; Zhang et al. 2015). Here, information with respect to underlying mechanisms of DA toxicity is limited. Moreover, epidemiological links between OP exposures, along with gene-environment interactions and PD support the need for increased mechanistic studies (Manthripragada et al. 2010). The present study advances understanding of DA neurotoxicity mechanisms using *C. elegans* model system, enzymatic activity assays, and computational approaches. CPF exposure in nematodes exhibited dose dependent loss of DA neurons with CEP and PDE neurons least and most vulnerable respectively. PDE neurons are late stage DA neurons (Wicks and Rankin 1996). In addition, CPF exposure led to a developmental delay. Our studies in L4 worms validated that vulnerability of PDE neurons was partially due to developmental delay (Supplementary information: S2, Table 1, Figure S2). Upon testing the effect on DA and ACh associated behavior in day 2 and day 3 washout experiments (worms were allowed to recover in absence of CPF), we observed that while the effect on ACh based behavior (accumulation of ACh culminating muscular paralysis) solely relies on presence of CPF, the effect on DA levels (delayed repulsion time owing to DA deficit) persists even after withdrawal, indicating that CPF induced DA neurotoxicity is likely irreversible at comparative doses relative to ACh effects. In comparison between ACh and DA associated behavior, we found that in two day studies ACh dependent paralysis surpassed the DA deficits, while the reverse was observed in the washout experiments owing to retention of DA deficits. Given that sporadic cases of PD are often associated to exposure to environmental factors such as pesticides, this finding relates to a plausible health risk resulting from pesticide exposures.

CPF is metabolized by CYP450 enzymes in mammals leading to formation of either CPF oxon and 3,5,6 trichloro-2-pyrindol (Tang et al. 2001). CPF oxon is most potent AchE inhibitor than its parent compound CPF (Wu and Laird 2003). Hence we tested the effect of CPF oxon Vs CPF in nematodes. Nematodes exposed to CPF oxon at equimolar concentrations exhibited relatively higher DA toxicity (EC_50_ of CPF oxon = 14.043 μM, CPF = 75.391 μM). Similarly, we observed significantly higher repulsion time (indicative of decreased DA levels) in response to CPF oxon.

Mitochondrial impairments and damage have been associated with Parkinson’s disease etiology (Exner et al. 2012; Foo et al. 2020; Keane et al. 2011). Specific pesticides, such as rotenone are known target mitochondrial function in all cells resulting in rather selective neurotoxicity of DA neurons due to specific sensitivity (Betarbet et al. 2000). Similar reports have also emerged for organophosphates (Karami-Mohajeri and Abdollahi 2013), indicating a plausible link between PD pathology and pesticides. Earlier reports have also shown deleterious effects of CPF on mitochondria (Singh et al. 2018; Toualbia et al. 2017; Turton et al. 2021). Hence, we tested the effect of CPF on the content of viable mitochondria viability, followed by testing of effect of CPF and CPF oxon on mitochondrial enzyme activity. We observed a significant reduction in content of viable mitochondria. Studies pertaining to mitochondrial function were furthered by directly measuring the effect of CPF and its active metabolite, CPF oxon on individual mitochondrial complex activities. While CPF oxon lacked any significant effect on complex activity, CPF exhibited a significant inhibition of Complex II, II+III and V. CPF oxon is considered highly neurotoxic and a key metabolite pertaining to CPF toxicity. Our results identified alternate mechanism, independent of the toxic metabolite CPF oxon, which presents considerable risk arising from repeated exposures. Interestingly, inhibitory effects on complex II and II+III were found to be reversed by Coenzyme Q1 (CoQ) supplementation. CoenzymeQ1 is an analog of CoQ10 (Aldrich), which acts as an electron carrier in mitochondrial electron transport chain (Lenaz et al. 2007), and as an antioxidant (Bentinger et al. 2007; Crane 2001). Notably, sheep dip poisoning which is associated with organophosphate toxicity exhibits symptoms similar to CoQ10 deficiency (Mantle and Hargreaves 2018). Our results also complemented the recent study conducted by Turton et al., 2021 pertaining to CoQ10 deficiency in response to CPF exposure and rescue of cell viability and Complex II+III activity upon CoQ10 administration (Turton et al. 2021). Notably while the studies conducted by Turton et al., 2021 were limited to citrate synthase activity and complex II+III activity, our studies assessed all five complexes. We also observed a significant inhibition of complex V in response to CPF. Although this inhibition was also possibly present in case of CPF oxon, but it was statistically insignificant. This effect could also be correlated with the symptoms such as muscle fatigue in individuals exposed to chlorpyrifos (Mantle and Hargreaves 2018). These findings identified mitochondrial physiology as a key target for DA neurotoxicity elicited by the parent compound. A clinical study on CPF metabolism conducted by Eyer et al., 2009 has highlighted presence of extreme variability (~100 fold) in conversion of CPF-to-CPF oxon in humans, rendering some individuals more vulnerable. This study also highlighted that within 0 to 4 hrs of CPF exposure 75% of CPF is converted to CPF oxon (Eyer et al. 2009). With respect to this, our results particularly highlight that while CPF oxon exerts neurotoxicity, nonmetabolized CPF is an inhibitor of mitochondrial complexes. Chronic CPF exposures might result in inhibition of mitochondrial complexes and hence pose a neurotoxic risk in the long term. Although there is extreme variability in metabolism of CPF, parent compound, CPF alone also induces neurotoxicity via secondary mechanisms such as mitochondrial affliction, which can modulate disease progression upon constant exposures.

Similar to the studies on CPF, we also tested the effect of CPF oxon on DA and ACh associated behavior through 2 day/48 hrs studies and washout experiments. A significant increase in repulsion time (indicative of depleted DA levels) and ACh mediated paralysis was observed in 1-nonanol assay and aldicarb assay, respectively. Conversely to what was observed in aldicarb assay with CPF, CPF exon retained its effect even in washout experiments, validating its highly potent effect on AchE (Wu and Laird 2003). Furthermore, a comparison between the effect on 1-nonanol assay and aldicarb assay indicated that the effect on DA associated behavior surpassed that of Ach mediated behavior.

We next tested the effect of low doses of CPF (in order to avoid developmental delay due to high CPF toxicity) on mRNA expression of genes related to organophosphate detoxification (*poml-1*/PON1), critical for organophosphate toxicity (*cyp-35A2* and *cyp-35A3*), related to apoptosis (*ced-4* and *ced-3*) and necrosis (*crt-1* and *vha-12*) in nematodes. The *poml-1 gene* is orthologous to human PON1 (Chen et al. 2016) (Chen et al. 2016). However, due to lack of evidence, it is still unclear that *poml-1* performs the same function as human PON1. *Cyp-35A2* and *cyp-35A3* are xenobiotic responsive genes in C. *elegans* and have been shown to upregulate upon CPF exposure (Roh et al. 2014). *Ced-4* and *ced-3* are associated with apoptosis in dopaminergic neurons, while *crt-1* and *vha-12* are linked to necrosis in nematodes (Pu and Le 2008). While there is no evidence of paraxonase activity in *C. elegans*, the notion behind testing the mRNA expression of *poml-1* was to ascertain, if *poml-1* could be a putative functional ortholog of PON1. A lack of any dose dependent effect on *poml-1* did not support this hypothesis. Next, we tested the effect on mRNA expression of genes, *cyp-35A2* and *cyp-35A3* which have been previously identified critical for organophosphate toxicity (Roh and Choi 2011; Roh et al. 2014). We observed a dose dependent increase in expression of these genes, corroborating the earlier findings with respect to the critical nature of these genes in OP toxicity (Roh and Choi 2011). Furthermore, this paved the way for mutant studies to confirm the role of these genes. Next, we tested the effect on genes related to cell death. While such genes are ~20 in C. *elegans*, only a few are expressed in DA neurons. Genes *ced-3* and *ced-4*, and *crt-1* and *vha-12*, are associated with apoptosis and necrosis in DA neurons (Pu and Le 2008). We observed only slight alterations in mRNA expression of genes *ced-3* and *crt-1*. qPCR data suggested a key role of *cyp-35A2* and *cyp-35A3*. Hence, we tested the effect of *cyp-35A2* and *cyp-35A3* mutations on DA cell loss. We found that both gene mutations result in amelioration of DA neurotoxicity. *Cyp-35A2* and *cyp-35A3* belong to *cyp-35a* family, besides being xenobiotic responsive genes, they are also involved in lipid metabolism in nematodes (Imanikia et al. 2015),(Menzel et al. 2007). Previous studies in *C. elegans* have shown upregulation of these genes in response to CPF exposure (Roh et al. 2014). Also, silencing has shown to lower gut lipid levels in *C. elegans* (Menzel et al. 2007). In addition, double mutants for *fat-5* and *cyp-35A2* have shown reduction in lipid content in *C. elegans* (Imanikia et al. 2015). Notably, CPF is known to accumulate in fat (Bakke et al. 1976; Eaton et al. 2008; Smith et al. 1967), due to prolonged half-life (Smith et al. 1967), which is further exacerbated by CPF induced accumulation of fat (Howell et al. 2016; Wang et al. 2021). *Cyp-35A2* and *cyp-35A3* are not homologous to human CPF metabolizing enzymes (i.e., CYP450 2B6). Thus, these mutations possibly results in fat reduction and hence curtailed CPF uptake/accumulation in *C. elegans*.

While the majority of CPF neurotoxicity is thought to arise from CPF oxon and resultant AchE inhibition by OP pesticides, our studies identify alternate mechanisms of parent compound neurotoxicity in dopaminergic neurons, where mitochondrial deficits and the role of fat modulatory genes, are particularly important. Figure 6 shows clustering of lipid metabolic assays such as the ER, PXR, PPAR gamma pathways, and the neurotoxic assay results. Figure 7A and 7B expands upon these assay results, grouping toxicity scores by gene targets in the first panel and neurotoxicity markers in the second. The result highlights the potential key events in the adverse outcome pathways of the parent OP compounds that result in DA neurotoxicity. Taken together, both the parent compound and oxon are likely to be neurotoxic to DA neurons and such neurotoxicity is reached through convergent, yet distinct mechanistic pathways. It is certainly possible that metabolic differences in humans influence the possible range of adverse neurological outcomes from CPF exposure. Although CPF is currently banned in U.S.A., the risk still perseveres owing to other organophosphates and constant use in other countries. This research will stimulate further interest in understanding the mechanistic aspects of organophosphate toxicity.

## 5. Funding

This work was supported by the National Institute of Environmental Health Sciences at the National Institutes of Health (R01ES025750 to JC; K99ES032488 to SRS) and the Ralph W. and Grace M. Showalter Research Trust (to JC).

## 6. Acknowledgements

The authors would like to thank Dr. Timothy Shafer and Dr. Kelly Carstens for their assistance on the ToxCast data analysis. The authors would also like to acknowledge C. elegans Gene Knockout Consortium. Strains were provided by the *Caenorhabdiditis* Genetics Center, funded by National Institute of Health Office of Research Infrastructure Programs (P40 OD010440).

## 7. Conflict of interest statement

The authors declare no conflicts of interest.

## Notes

### Competing Interest Statement

The authors have declared no competing interest.

