## Supplemental data for "Complementary biological and computational approaches identify distinct mechanisms of chlorpyrifos versus chlorpyrifos oxon induced dopaminergic neurotoxicity"

**Supplementary Information:**

**S1. Methods**

***1.1. Genetic Cross to study the effect of cyp-35A2 and cyp-35A3 mutation on PFOS-mediated neurodegeneration***: A genetic cross between BZ555 (dat-$1_p$::GFP) and VC710 (*cyp-35A2*(gk317)), RB2046 (*cyp-35A3(*ok2709) was conducted as previously described (Pu and Le 2008), with slight modifications. Briefly, BZ555 males were crossed with VC710, and RB2046 hermaphrodites. F-1 progeny showing GFP labeled neurons were transferred to NGM–OP50 plates (35 mm plates). Worms (F2) picked from plates with 100% positive worms were genotyped post egg-laying. Worms with homozygous deletions (*cyp-35A2* and *cyp-35A3,* designated as BZVC710 and BZRB2046) were grown and used for the studies.

*Table 1: Sequence of the primers used*

| | Gene Name | Primer Sequence 5′ to 3′ |
|---|---|---|
| 1 | ***gpd-1*** **F** | AAA GTC ATT CCG GAG CTG AAC GGA |
| | ***gpd-1*** **R** | AGC GGC CTT GAC TAC CTT CTT GAT |
| 2 | ***poml-1*** **F** | CCT GGA GAA TGT GGC GTT AT |
| | ***poml-1*** **R** | CGG TAC GAT TGT CGG AGT AAA |
| 3 | ***ced-3*** **F** | GAC GGA GTT CCT GCA TTT CT |
| | ***ced-3*** **R** | CTT GGC TCG GCT TCT TTC T |
| 4 | ***ced-4*** **F** | GAT GCC TCC TGG AGT TGA TAT AC |
| | ***ced-4*** **R** | TGA GAA GAG CTC CAC GTT TG |
| 5 | ***crt-1*** **F** | CAT CCT CAA CTC CGA CAA TAC C |
| | ***crt-1*** **R** | CCT CTG GCT TCT TTG CAT CT |
| 6 | ***vha-12*** **F** | CCA TCC CTT TCC CGA CTT ATG |
| | ***vha-12*** **R** | CGA TAG CGT AAC AAG CGT AGA G |

**S2. Results:**

**Supplementary Figure 1**

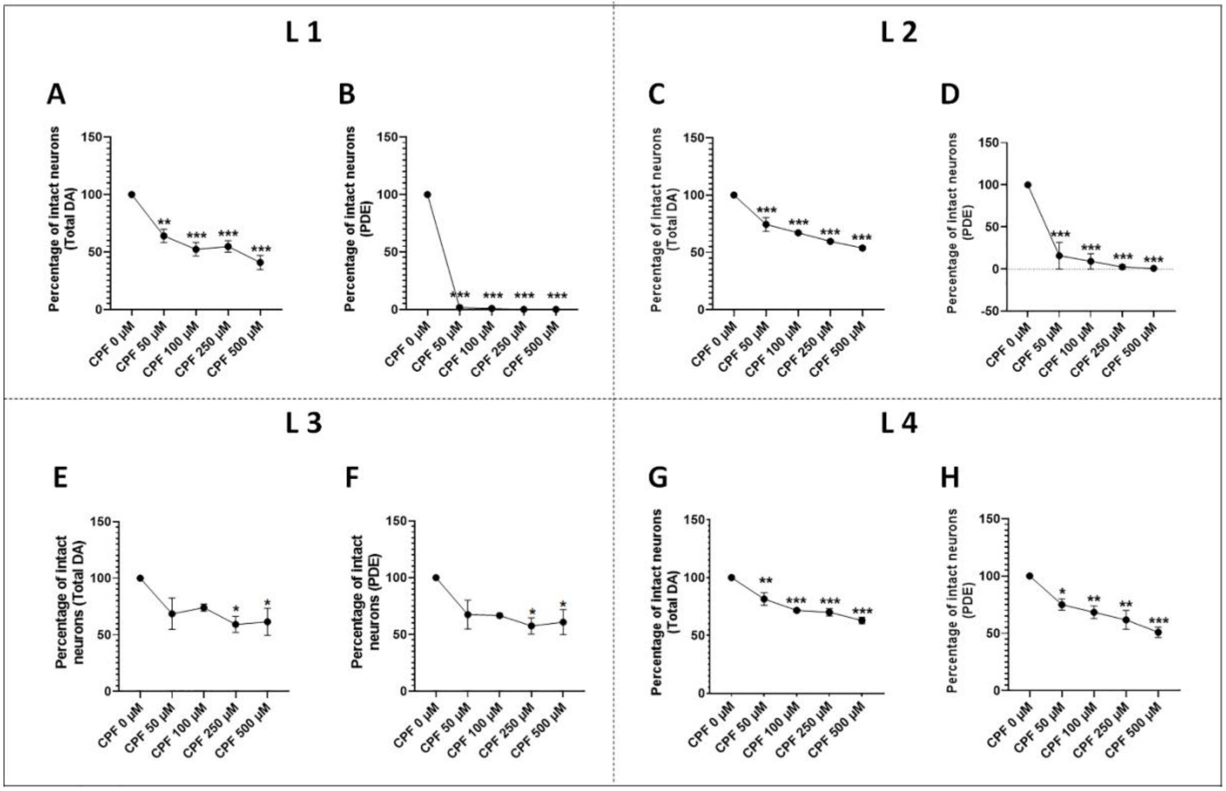


***Supplementary Figure 1. CPF neurotoxicity to PDE neurons is molting stage dependent***. Data in section 3.1, suggest that vulnerability of the PDE neurons could be partly due to the developmental delay. Therefore, we treated the worms at different molting stages (L1, L2, L3, and L4) to test this hypothesis. As expected, we found that the number of live PDE neurons was impacted due to the developmental delay. The percentage of intact neurons for both total neurons and PDE neurons has been shown in table 1. Overall our data indicated that PDE neurotoxicity is apparently higher in L1 treated worms partly due to the developmental delay and in part due to toxic effects of CPF. This effect diminishes from L2 to L4 treated worms (Figure S1 ***B***, ***D***, ***F***, ***H***). Comparing the effect of CPF on total neurons, it can also be concluded that the early-stage neurons are more susceptible to CPF mediated neurotoxicity (***Figure S1 A***, ***C***, ***E***, ***G***). Specific experimental details: CPF DA cell loss was assessed after 72 hrs treatments at different molting stages for different sub class of neurons, CEP, ADE, PDE. Worms treated at L1 exhibited dose dependent loss of DA neurons, Total (***A***), and PDE (***B***). Worms treated at L2 exhibited dose dependent loss of DA neurons, total (***C***), and PDE (***D***). Worms treated at L3 exhibited dose dependent loss of DA neurons, Total (***E***), and PDE (***F***). Worms treated at L4 exhibited dose dependent loss of DA neurons, Total (***G***), and PDE (***H***). Data are presented as mean ± S.E.M. Data were analyzed using one-way ANOVA followed by Dunnett's post hoc test. *$p<.05$, **$p<.005$, and ***$p<.001$ (n =3).

*Supplementary Table 1. Quantification of the effects of CPF on Total and PDE neurons in worms treated at different molting stages.*

| | Concentration of CPF | Percentage of intact neurons (Total) | *p*-value | Percentage of intact neurons (PDE) | *p*-value |
|---|---|---|---|---|---|
| **L1** | 0 μM | 100 ± 0.000 | | 100 ± 0.000 | |
| | 50 μM | 63.958 ± 5.867 | =0.0021 | 1.667 ± 1.667 | <0.0001 |
| | 100 μM | 52.292 ± 5.977 | =0.0003 | 0.833 ± 0.833 | <0.0001 |
| | 250 μM | 54.792 ± 5.158 | =0.0004 | 0.000 ± 0.000 | <0.0001 |
| | 500 μM | 40.833 ± 6.106 | <0.0001 | 0.000 ± 0.000 | <0.0001 |
| **L2** | 0 μM | 100 ± 0.000 | | 100 ± 0.000 | |
| | 50 μM | 74.375 ± 6.006 | =0.0003 | 15.833 ± 15.833 | =0.0001 |
| | 100 μM | 67.083 ± 1.102 | <0.0001 | 9.167 ± 9.167 | <0.0001 |
| | 250 μM | 59.583 ± 1.267 | <0.0001 | 2.500 ± 2.500 | <0.0001 |
| | 500 μM | 53.750 ± 1.654 | <0.0001 | 0.833 ± 0.833 | <0.0001 |
| **L3** | 0 μM | 100 ± 0.000 | | 100 ± 0.000 | |
| | 50 μM | 63.958 ± 5.867 | =0.002 | 67.500 ± 12.829 | <0.000 |
| | 100 μM | 52.292 ± 5.977 | =0.003 | 66.667 ± 1.667 | <0.000 |
| | 250 μM | 54.792 ± 5.158 | =0.000 | 57.500 ± 7.217 | <0.000 |
| | 500 μM | 40.833 ± 6.106 | <0.000 | 60.833 ± 11.024 | <0.000 |
| **L4** | 0 μM | 100 ± 0.000 | | 100 ± 0.000 | |
| | 50 μM | 74.375 ± 6.006 | =0.000 | 75.000 ± 5.000 | =0.000 |
| | 100 μM | 67.083 ± 1.102 | <0.000 | 68.333 ± 5.465 | <0.000 |
| | 250 μM | 59.583 ± 1.267 | <0.000 | 61.667 ± 8.207 | <0.000 |
| | 500 μM | 53.750 ± 1.654 | <0.000 | 50.833 ± 4.410 | <0.000 |

**Supplementary Figure 2**

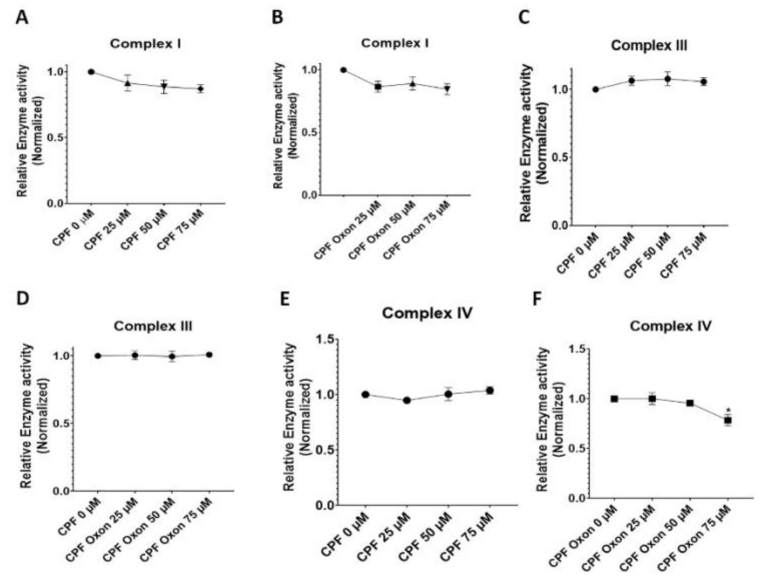


***Supplementary Figure S2. Pilot study on mitochondrial respiration targets.*** In order to identify if the CPF and its metabolite CPF Oxon have any effect on mitochondrial complex enzyme activity (EA), we performed assays for assessment of EA in mitochondria isolated from rat liver. Initial studies were conducted on three doses (25 to 75 μM). Subsequently, detailed investigation using a broader range was conducted where the effect was observed. ***CPF and CPF Oxon do not affect Complex I activity****:* A significant number of studies have linked complex I inhibiton to PD pathology (Gonzalez-Rodriguez et al. 2021; Keeney et al. 2006; Subrahmanian and LaVoie 2021). Hence, we studied the effect of CPF and CPF Oxon on complex I EA. We did not observe any significant alteration in complex I EA in mitochondria treated with CPF and CPF Oxon (***Figure*** S2 ***A, B***). ***CPF and CPF Oxon do not affect Complex III activity****:* Further we also tested CPF and CPF oxon effects on complex III EA. We did not observe any significant change in complex III activity in mitochondria treated with CPF and CPF Oxon (***Figure S2 C, D***).***CPF and CPF Oxon do not affect Complex IV activity****:* Next, we tested the effect of CPF and CPF Oxon on complex IV EA. CPF treatment did not affect complex IV EA (***Figure S2 E***). However, CPF Oxon treatment did show some effect, but only at the highest dose, 75 μM (0.784 ± 0.057, $p = 0.0194$) in comparison to control (1.000 ± 0.000) (***Figure S2 F***). Data analyzed using one-way ANOVA followed by Dunnett's post hoc test. *p<.05, **p<.005, and ***p<.001 (n =3). Data were analyzed using one-way ANOVA followed by Dunnett's post hoc test. *p<.05(n =3).

**Supplementary Figure 3**

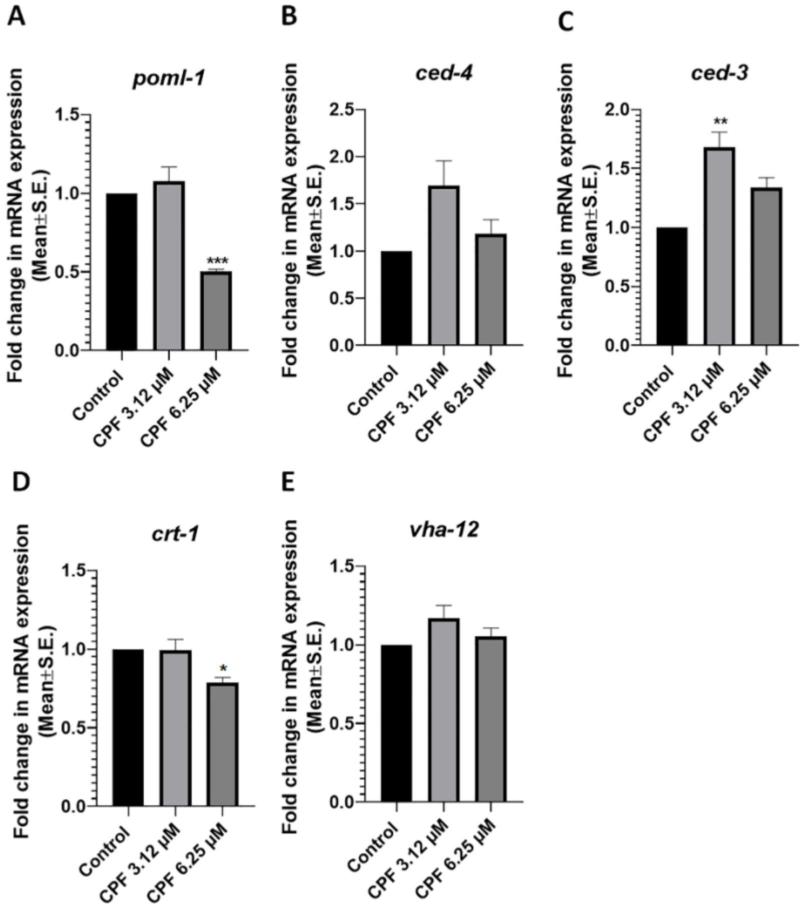


***Supplementary Figure 3. Effect of CPF on expression of genes related to PON1 ortholog, apoptosis and necrosis***. After determining the neurotoxic effects of CPF on DA neurons, we studied the effect of CPF on expression of genes related to PON1 ortholog (*poml-1),* apoptosis (*ced-3* and *ced-4*), and necrosis (*crt-1* and *vha-12*). The experiments were conducted on worms exposed at L1 stages. Lower doses of CPF were studied to minimize any confounding factors arising from toxicity of CPF. We observed a significant decrease in mRNA expression of *poml-1* at CPF 6.25 µM (0.500 ± 0.017, $p = 0.0009$) in comparison to control (1.000 ± 0.000) (***Figure S3 A***). In case of genes related to apoptosis (*ced-3* and *ced-4*), a significant increase in expression of *ced-3* at CPF 3.12 µM (1.680 ± 0.127, $p = 0.0029$) in comparison to control (1.000 ± 0.000) (***Figure S3 B***). Any effect on mRNA expression of *ced-4* was statistically insignificant (***Figure S3 C***). In case of genes related to necrosis (*crt-1* and *vha-12*), a slight, yet significant decrease in expression of *crt-1* at CPF 6.25 µM (0.786 ± 0.033, $p = 0.0235$) in comparison to control (1.000 ± 0.000) (***Figure S3 D***). Any effect on mRNA expression of *vha-12* was statistically insignificant **(Figure S3 E)**. Data are presented as mean ± S.E.M. Data were analyzed using one-way ANOVA followed by Dunnett's post hoc test. *p<.05, **p<.005, and ***p<.001 (n =3).
